# Modulating D1 rather than D2 receptor-expressing spiny-projection neurons corresponds to optimal antipsychotic effect

**DOI:** 10.1101/2021.08.03.454992

**Authors:** Seongsik Yun, Ben Yang, Madison M. Martin, Nai-Hsing Yeh, Anis Contractor, Jones G. Parker

## Abstract

Overactive dopamine transmission in psychosis is predicted to unbalance striatal output via D1- and D2-dopamine receptor-expressing spiny-projection neurons (SPNs). Antipsychotic drugs are thought to re-balance this output by blocking D2-receptor signaling. Here we imaged D1- and D2-SPN Ca^2+^ dynamics in mice to determine the neural signatures of antipsychotic effect. Initially we compared effective (clozapine and haloperidol) antipsychotics to a candidate drug that failed in clinical trials (MP-10). Clozapine and haloperidol normalized hyperdopaminergic D1-SPN dynamics, while MP-10 only normalized D2-SPN activity. Clozapine, haloperidol or chemogenetic manipulations of D1-SPNs also normalized sensorimotor gating. Given the surprising correlation between clinical efficacy and D1-SPN modulation, we evaluated compounds that selectively target D1-SPNs. D1R partial agonism, antagonism, or positive M4 cholinergic receptor modulation all normalized the levels of D1-SPN activity, locomotion, and sensorimotor gating. Our results suggest that D1-SPN activity is a more relevant therapeutic target than D2-SPN activity for the development of effective antipsychotics.

## Main

Antipsychotic drugs have been used to manage the symptoms of psychotic disorders for over half a century. Very early on, it was recognized that excess dopamine might contribute to psychosis^1^, and that antipsychotic drugs may act on the dopamine system^2^. A close association between D2- like dopamine receptor binding and antipsychotic effect bolstered this idea^3^ and a dopamine hypothesis for psychotic disorders like schizophrenia or the mechanistic basis of the drugs for these disorders^4^. Since that time, intense therapeutic development efforts have sought to further fine tune D2-like receptor signaling. These efforts yielded compounds with lower D2 receptor (D2R) binding affinities^5^, selectivity for specific D2-like receptors^6^, partial agonists that ‘stabilize’ D2R signaling^7^, functionally selective D2R ligands^8^, and compounds that target signaling pathways downstream from D2Rs^9^. Despite these remarkable pharmacological advances, comparatively little progress has been made in terms of the real-world efficacy of antipsychotic treatments. Given this discrepancy, there is an immediate need to understand the effects of these drugs on the function of intact neural circuits that are thought to underlie psychosis.

In schizophrenia, increased dopamine transmission is thought to imbalance the rates of activity in the striatum’s principal output neurons, the D1R- and D2R-expressing spiny projection neurons (SPNs)^10^. Specifically, activation of Gα_s_-coupled D1Rs and Gα_i_-coupled D2Rs is predicted to increase D1- and decrease D2-SPN activity^11^. D1- and D2-SPNs input to the direct and indirect basal ganglia pathways, respectively, which converge to modulate basal ganglia output to the thalamus. In theory, treatments that normalize the activity of either or both SPN types could normalize basal ganglia output. However, the receptor pharmacology of antipsychotic drugs predicts that they preferentially normalize D2-SPN activity. However, whether increased dopamine unbalances D1- and D2-SPN activity and whether antipsychotic drugs normalize this imbalance through selective effects on D2-SPNs has never been directly tested *in vivo*.

Using a miniature microscope to image D1- and D2-SPN Ca^2+^ activity *in vivo*, we and others showed that D1- and D2-SPNs co-activate in spatially clustered ensembles and scale their levels of activity with locomotor speed in a balanced manner^12, 13^. Conditions modeling parkinsonism and dyskinesia disrupt both the levels and spatially clustered dynamics of D1- and D2- SPN activity^12^. Importantly, the extent to which mainstay or candidate treatments for Parkinson’s disease normalize these dynamics is more predictive real-world efficacy than behavioral measures in an animal model of parkinsonism^12^. Given the great number of neurological and psychiatric diseases for which striatal dysfunction is implicated^14–20^, the ability to examine how these dynamics are disrupted in other disease states and normalized by their treatments is a powerful tool for understanding brain pathophysiology and therapeutic effect.

In the present study, we recorded D1- and D2-SPN Ca^2+^ activity in the dorsomedial striatum (DMS) to determine how antipsychotic drugs modulate their dynamics under normal and hyperdopaminergic conditions. Initially we sought to use this readout to determine whether we could distinguish between antipsychotic drugs or drug candidates with varying clinical efficacies and side-effect profiles. Specifically, we compared two effective antipsychotics (clozapine and haloperidol) and one ineffective antipsychotic drug candidate (MP-10)^9, 21^. Presently, the different efficacies and side-effect propensities of antipsychotic drugs are best understood by their different brain receptor binding profiles^22^. For instance, D2R binding is thought to underlie the antipsychotic effects of clozapine and haloperidol. However, haloperidol’s greater selectivity and binding affinity for D2Rs is thought to underlie its greater propensity for motor side effects like parkinsonism and dyskinesia. Likewise, clozapine’s greater affinity for serotonin receptors may underlie its superior antipsychotic efficacy.

Although this taxonomical approach is extremely useful, D2 and the other receptors bound by antipsychotic drugs are widely distributed throughout the brain, making it difficult to link a specific drug’s receptor interactions to its specific therapeutic profile. MP-10 exemplifies the limitations of linking specific disease symptoms to receptor signaling pathways in this way. MP-10 inhibits PDE10A, a striatally enriched enzyme whose inhibition increases the levels of the second-messenger cAMP in the striatum^23^. Because D2R signaling inhibits cAMP production^11^, MP-10 effectively recapitulates D2R antagonism with specificity for the striatum. Given the strong linkage between striatal D2R binding and antipsychotic effect, MP-10 was predicted to be antipsychotic, with possibly fewer of the adverse effects associated with brain-wide D2R antagonism. Although this prediction is logical within a receptor-symptom conceptual framework, MP-10 had no antipsychotic effect in patients with schizophrenia^9^.

In terms of behavior, clozapine, haloperidol, and MP-10 all suppressed basal locomotion and attenuated hyperlocomotion following treatment with the dopamine releaser amphetamine. A drug’s ability to suppress of amphetamine-driven locomotion in rodents is a common indicator of antipsychotic potential. However, not every drug that attenuates amphetamine-driven locomotion also has antipsychotic activity. MP-10 is one of many such examples that underscore the limited predictive value of this assay, particularly when behavior is the primary readout. In terms of neural activity, we found that amphetamine treatment increased D1- and decreased D2-SPN activity levels, and differentially altered their spatiotemporal dynamics. Despite their similar effects on locomotion, clozapine, haloperidol and MP-10 each had distinct effects on these hyperdopaminergic ensemble dynamics. Surprisingly, the selective normalization of D1-, rather than D2-SPN dynamics was associated with clinical antipsychotic effect, and the ineffective drug candidate (MP-10) actually exacerbated amphetamine’s effects on D1-SPN activity. Thus, by examining the neural, rather than behavioral correlates of antipsychotic drug effect, we could retrospectively distinguish between three drugs known to have different clinical efficacies.

Given the correlation between D1-SPN modulation and clinical antipsychotic effect, we asked whether D1-SPN modulation was sufficient to normalize amphetamine-driven changes in behavior. Chemogenetic inhibition of D1-SPNs in the DMS was sufficient to normalize amphet-amine-driven hyperlocomotion and deficits in sensorimotor gating, another common behavioral measures of antipsychotic drug potential. Next we tested whether compounds targeting receptors enriched in D1-SPNs might be therapeutically active for dopamine-driven psychosis. Three compounds targeted to either D1Rs (SKF38393 and SCH23390) or M4 cholinergic receptors (VU0467154)^24^ all normalized hyperdopaminergic D1-SPN dynamics and behavioral measures of antipsychotic drug potential.

Taken together, our results highlight the power of a neural ensemble imaging approach for distinguishing between treatments for brain diseases like psychosis and for uncovering the mechanistic basis for their efficacy. This approach has uncovered the surprising finding that targeting D1-SPNs may provide greater therapeutic benefit than traditional D2R-based antipsychotic treatments. This new perspective and its underlying technical advances provide a framework for developing novel and potentially more comprehensive treatments for psychosis.

## Results

### D1- and D2-SPN dynamics under normal and hyperdopaminergic conditions

To record D1- or D2-SPN activity *in vivo*, we used a virus to conditionally express the fluorescent Ca^2+^ indicator GCaMP7f in the DMS of *Drd1a^Cre^* (D1-Cre) or *Adora2a^Cre^* (A2A-Cre) mice, respectively. We implanted an optical guide tube and microendoscope into the DMS and mounted the mice with a miniature fluorescence microscope (**Fig. 1a** and **Extended Data Fig. 1a**). This approach allowed us to monitor Ca^2+^ activity in hundreds of individual D1- or D2- SPNs as mice freely explored an open field arena (**Fig. 1b**; 233 ± 11 D1-SPNs over 189 imaging sessions and 161 ± 10 D2-SPNs over 172 sessions; mean ± s.e.m). D1- and D2-SPNs had similar event rates and similarly increased their levels of activity with locomotor speed (**Fig. 1c** “Vehicle” and **Extended Data Fig. 1b**). Under control conditions, Ca^2+^ event rates were 0.5 ± 0.05 events·min^-1^ for D1-SPNs and 0.6 ± 0.08 events·min^-1^ for D2-SPN during periods of rest (locomotor speed < 0.5 cm·s^-1^). During periods of movement (speed >= 0.5 cm·s^-1^) D1-SPNs had rates of 1.6 ± 0.07 events·min^-1^ and D2-SPNs had rates of 1.8 ± 0.1 events·min^-1^ (*P* = 0.5 for rest and *P* = 0.2 for movement; *N* = 11 D1-Cre and 10 A2A-Cre mice; Wilcoxon rank-sum test). Consistent with previous findings, D1- and D2-SPNs both exhibited spatiotemporally coordinated patterns of activity, whereby proximal pairs of cells (separated 25–125 µm) had more temporally overlapped Ca^2+^ events than distal cell pairs (**Extended Data Fig. 1c**)^12^. In contrast to the overall levels of D1- and D2-SPN activity, this co-activity among proximal cell pairs decreased with increased locomotor speed (**Extended Data Fig. 1d**). To account for the relationship between locomotor speed and these parameters of D1- and D2-SPN activity, we performed all subsequent analyses as a function of each mouse’s running speed.

**Fig. 1:**
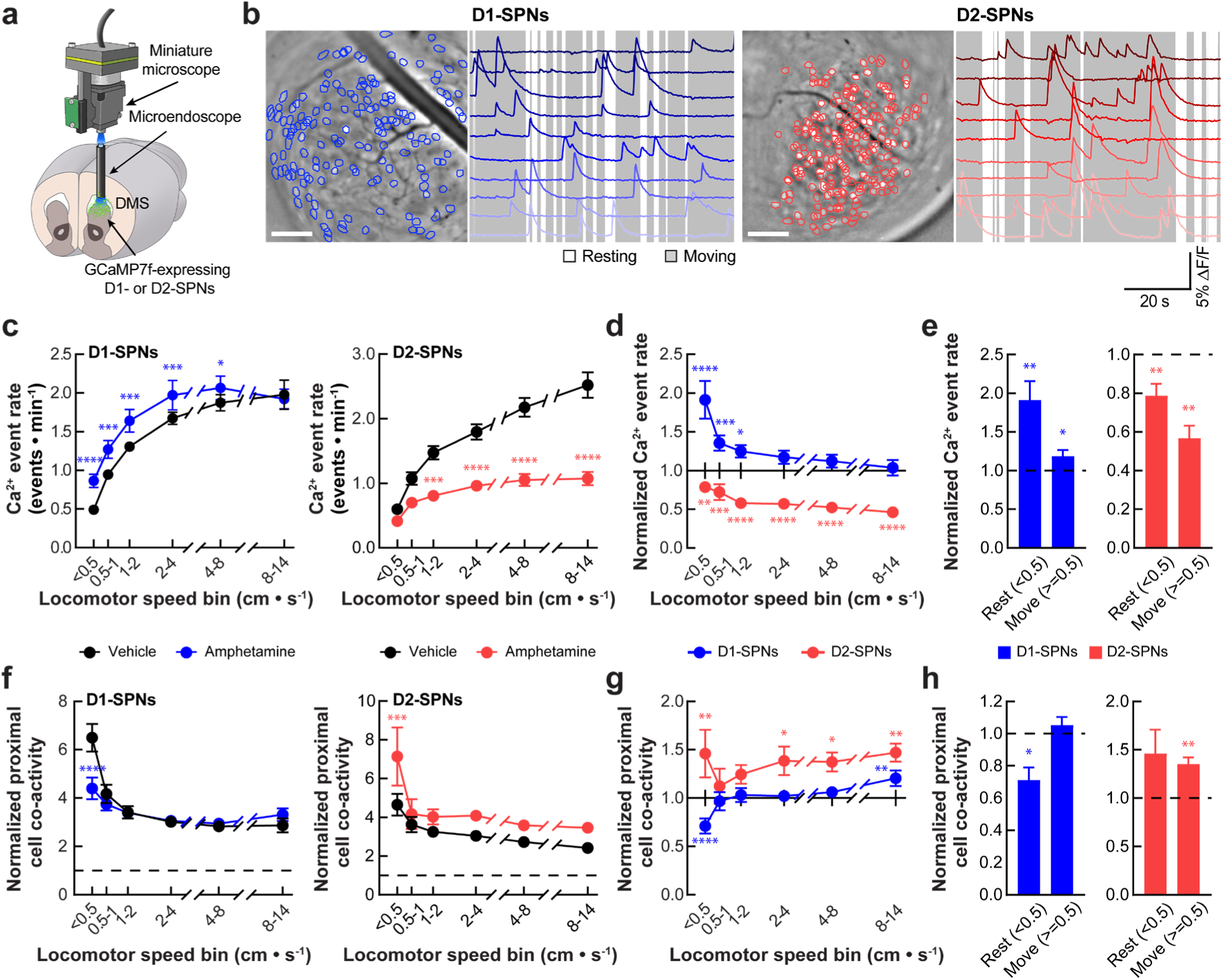
Effects of amphetamine treatment on D1- and D2-SPN Ca^2+^ activity in freely behaving mice. **a**, We used a miniature microscope and microendoscope to image Ca^2+^ activity in D1- and D2-SPNs by expressing GCaMP7f in the dorsomedial striatum. **b**, Cell centroid locations overlaid on the mean fluorescence images of dorsomedial striatum and example Ca^2+^ activity traces from D1-SPNs and D2-SPNs in representative D1- (*left*) and A2A-Cre (*right*) mice. Scale bar: 100 µm. **c,** Effects of vehicle or amphetamine on Ca^2+^ event rates in D1- and D2-SPNs across increasing locomotor speed bins. **d**, Ca^2+^ event rates in D1- and D2-SPNs following amphetamine treatment, normalized values following vehicle only treatment across different speed bins. **e**, Effects of amphetamine on Ca^2+^ event rates in D1- and D2-SPN during resting (< 0.5 cm·s^-1^) and moving (>= 0.5 cm·s^-1^) speed bins, normalized to mean values following vehicle only treatment. **f**, Co-activity of proximal D1- and D2-SPN pairs (25–125 µm separation) across different speed bins, normalized to temporally shuffled comparisons following vehicle or amphetamine treatment. **g**, Co-activity of proximal D1- and D2-SPN pairs at different speed bins after amphetamine treatment, first normalized to temporally shuffled comparisons and then to the mean, shuffle-normalized values following vehicle only treatment. **h**, Effects of amphetamine on co-activity of proximal D1- and D2-SPN pairs during resting and moving speed bins, normalized to temporally shuffled comparisons and then to values observed following vehicle only treatment. Data are expressed as mean ± s.e.m. (*N* = 11 D1-Cre and *N* = 10 A2A-Cre mice; \*\*\*\**P* < 0.0001, \*\*\**P* < 0.001, \*\**P* < 0.01 and \**P* < 0.05 comparing amphetamine to vehicle treatment; Holm-Sidak’s multiple comparison test for **c**, **d**, **f**, and **g**; Wilcoxon signed-rank test for **e** and **h**).

To determine how the increased striatal dopamine release in diseases such as psychosis, may affect these D1- and D2-SPN ensemble dynamics, we treated mice with amphetamine, which induces an efflux of cytoplasmic dopamine through dopamine transporter^25^. Consistent with excitatory D1R and inhibitory D2R activation, amphetamine treatment (2.5 mg·kg^-1^) increased D1- and decreased D2-SPN activity levels (**Fig. 1c**). These effects were dependent on locomotor speed, with greater D1-SPN activation during periods of rest and more D2-SPN suppression during movement (**Fig. 1c–e**). Amphetamine treatment also differentially altered the spatiotemporal dynamics of D1- and D2-SPNs in a speed-dependent manner (**Fig. 1f**). Amphetamine disruption of proximal D1-SPN co-activity and augmentation of proximal D2-SPN co-activity were most pronounced during periods of rest and movement, respectively (**Fig. 1f–h**). Taken together, amphetamine treatment diametrically altered the levels and spatiotemporal dynamics of D1- and D2-SPN ensembles.

### The neural ensemble correlates of antipsychotic drug efficacy

Next we asked whether we could use these dynamics to distinguish between three antipsychotic drugs with different clinical efficacies and side-effect profiles. We compared clozapine, a highly efficacious antipsychotic with few motor side effects, to haloperidol, a moderately efficacious antipsychotic with a high motor side effect propensity, and MP-10, an antipsychotic drug candidate that recently failed in a clinical trial for schizophrenia^26, 27^.

We monitored behavior and recorded D1- or D2-SPN Ca^2+^ activity following treatment with vehicle or a low/high dose of each drug followed by amphetamine (**Fig. 2a**). Under normal conditions, during 15-min before amphetamine treatment, both high and low doses of all three drugs inhibited locomotor activity (**Fig. 2b** and **Extended Data Fig. 2a**). Despite their similar effects on locomotion under normal conditions, the profiles of each drug’s effects on D1- and D2-SPN activity levels differed. Clozapine treatment selectively increased D1-SPN activity, haloperidol increased both D1- and D2-SPN activity at the higher dose, while MP-10 only increased D2-SPN activity (**Fig. 2c, d**). By contrast, all three drugs had similarly negligible effects on the degree of spatiotemporally coordinated D1- and D2-SPN activity at either dose (**Extended Data Fig. 3a, b**).

**Fig. 2:**
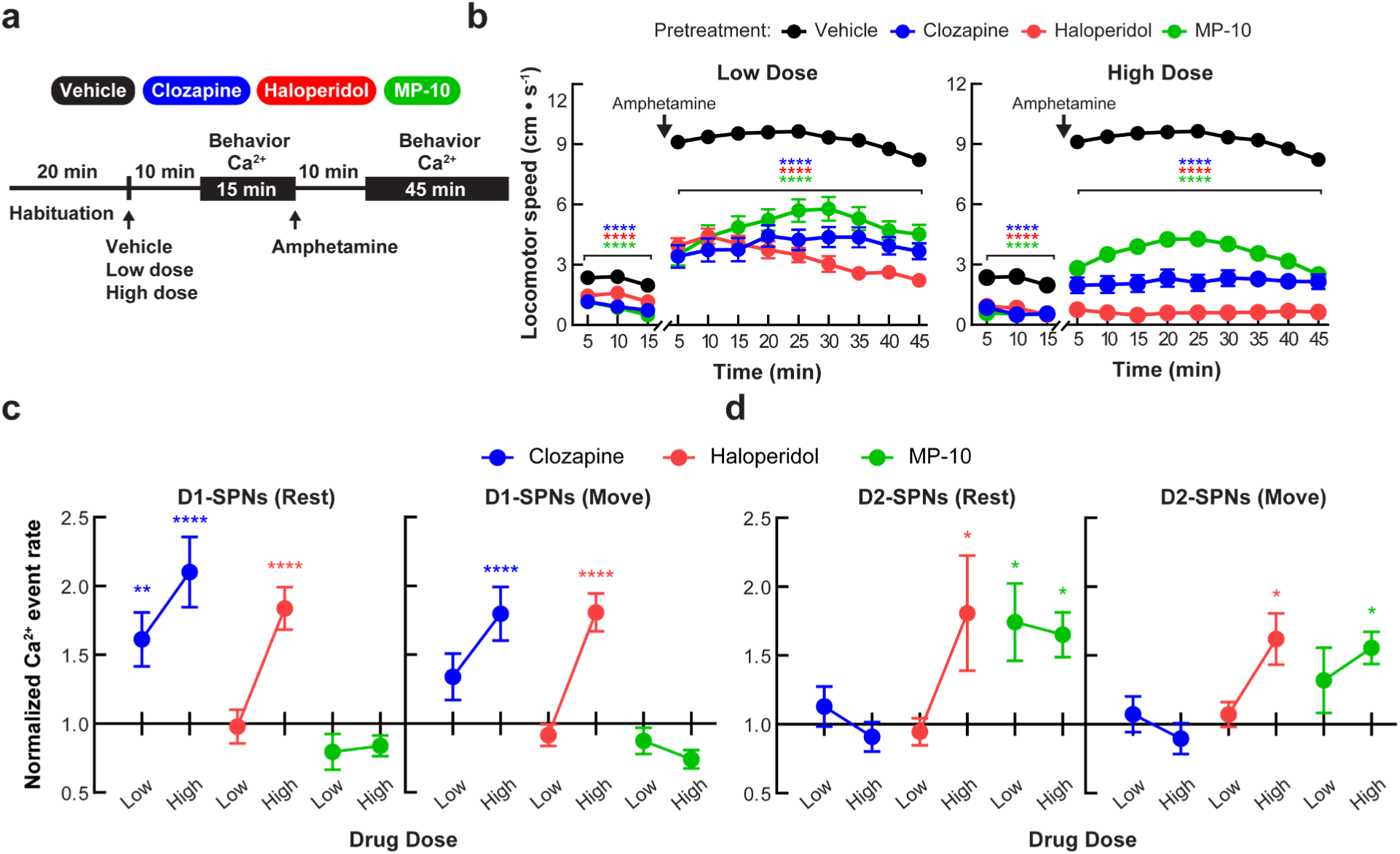
Effects of antipsychotic drugs on normal D1- and D2-SPN activity levels and spontaneous or amphetamine-driven locomotion. **a**, To record behavior and Ca^2+^ activity, we habituated the mice to the open field arena for 20 min before drug injection. After administering vehicle or a dose of antipsychotic drug, we recorded behavior and Ca^2+^ activity for 15 min, administered amphetamine, and recorded Ca^2+^ activity for an additional 45 min. All recordings began 10 min after vehicle, antipsychotic, or amphetamine treatment. We administered different antipsychotic drug doses on consecutive days and gave the mice a day off between the different drugs. **b**, Locomotor activity during the first 15 min recording period following vehicle or antipsychotic drug treatment and the 45 min recording period after amphetamine treatment (*N* = 31 mice; \*\*\*\**P* < 0.0001 comparing drug to vehicle and drug + amphetamine to vehicle + amphetamine treatment; Holm-Sidak’s multiple comparison test). **c**, **d**, Ca^2+^ event rates in D1-SPNs (**c**) and D2-SPNs (**d**) after low or high dose of antipsychotic drug treatment during rest and movement, normalized to event rates following vehicle only treatment. Data are expressed as mean ± s.e.m. (*N* = 11 D1-Cre and *N* = 10 for A2A-Cre mice; \*\*\*\**P* < 0.0001, **P < 0.01 and *P < 0.05 comparing drug dose to vehicle treatment; Holm-Sidak’s multiple comparison test).

**Fig. 3:**
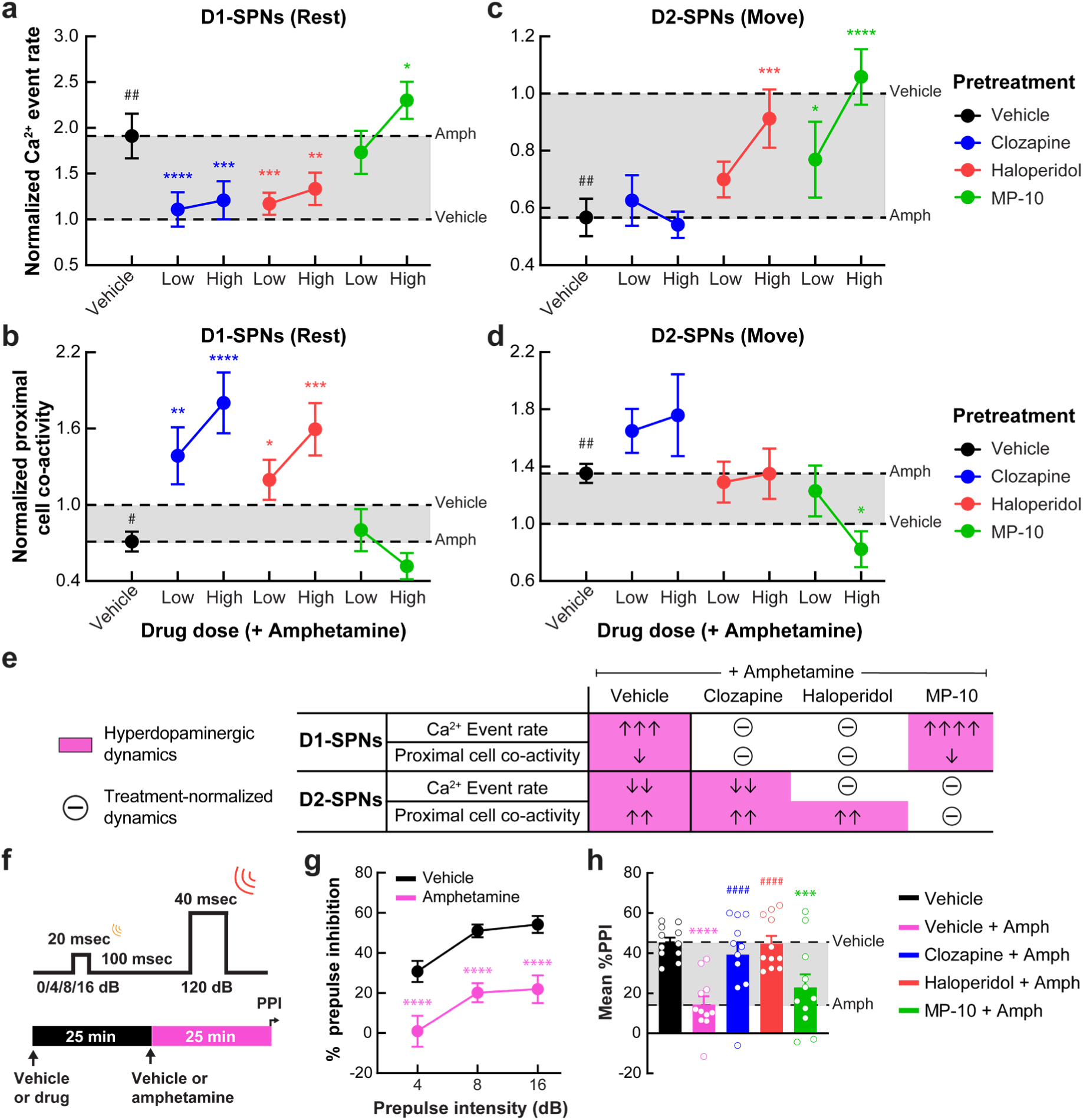
Differential effects of antipsychotic drugs on D1- and D2-SPN dynamics and sensorimotor gating under hyperdopaminergic conditions. **a**, **b**, Ca^2+^ event rates (**a**) and proximal co-activity (**b**) of D1-SPNs during periods of rest (locomotor speed < 0.5 cm·s^-1^) following vehicle or drug + amphetamine treatment, normalized to values following vehicle only treatment. **c**, **d**, Ca^2+^ event rates (**c**) and proximal co-activity of D2-SPNs (**d**) during periods of movement (locomotor speed >= 0.5 cm·s^-1^) following vehicle or drug + amphetamine treatment, normalized to values following vehicle only treatment (*N* = 11 D1-Cre and *N* = 10 A2A-Cre mice; ^#^*P* < 0.05 and ^##^*P* < 0.01 comparing amphetamine to vehicle treatment; \*\*\*\**P* < 0.0001, \*\*\**P* < 0.001, \*\**P* < 0.01 and \**P* < 0.05 comparing drug + amphetamine to vehicle + amphetamine treatment; Holm-Sidak’s multiple comparison test). **e**, Summary of the effects of the different antipsychotic drugs on amphetamine-disrupted D1- and D2-SPN ensemble dynamics. Pink shading indicates the hyperdopaminergic neural ensemble dynamics. Encircled “–” denotes antipsychotic-normalized changes. Arrows denote effect sizes compared to vehicle only (for vehicle + amphetamine) or to vehicle + amphetamine (for antipsychotic + amphetamine; one arrow >= 10%, two arrows >= 30%, and three arrows >= 50% statistically significant effect sizes; four arrows denotes the exacerbation of D1-SPN hyperactivity by MP-10 pre-treatment). **f**, We injected vehicle or antipsychotic drug 25 min before amphetamine treatment and started measuring PPI 25 min after amphetamine treatment. **g**, **h**, Percent PPI of startle response at 4, 8, and 16 dB pre-pulse intensities following vehicle or amphetamine only treatment (**g**) and mean percent PPI across all pre-pulse intensities following vehicle or drug + amphetamine treatment (**h**). All data are expressed as mean ± s.e.m. (*N* = 11; \*\*\*\**P* < 0.0001, \*\*\**P* < 0.001 compared to vehicle only treatment; ^####^P < 0.0001 comparing drug + amphetamine to vehicle + amphetamine treatment; Holm-Sidak’s multiple comparison test).

Under hyperdopaminergic conditions, both doses of all three drugs diminished ampheta-mine-driven hyperlocomotion (**Fig. 2b** and **Extended Data Fig. 2a**). Despite their comparable effects on behavior, the drugs differentially reversed the altered levels and spatiotemporal dynamics of D1- and D2-SPN activity. Both clozapine and haloperidol normalized the spatiotemporally de-correlated D1-SPN hyperactivity after amphetamine, while MP-10 exacerbated D1-SPN hyperactivity and had no effects on proximal D1-SPN co-activity (**Fig. 3a, b**). By comparison, MP-10 completely normalized the hyper-correlated D2-SPN hypo-activity with amphetamine, while haloperidol selectively normalized D2-SPN hypoactivity, and clozapine had no effects on the hyperdopaminergic ensemble dynamics of D2-SPNs (**Fig. 3c, d**). Taken together, our results show that the two clinically efficacious drugs normalized D1-SPN dynamics, while the inefficacious drug MP-10 only normalized D2-SPN dynamics under hyperdopaminergic conditions (**Fig. 3e**). Moreover, the most clinically efficacious drug clozapine exclusively normalized D1-SPN activity.

Given their disparate effects on D1- and D2-SPN activity but equivalent effects on amphetamine-driven locomotion, we next asked whether another behavioral assay might differentiate these three drugs treatment. In addition normalizing amphetamine-driven hyperlocomotion, antipsychotic drugs also normalize the disruption of sensorimotor gating as measured in rodents by pre-pulse inhibition (PPI)^28^. We pretreated mice with vehicle or a high dose of each antipsychotic drug followed by vehicle or amphetamine and measured PPI (**Fig. 3f**). Amphetamine treatment disrupted PPI at all pre-pulse intensities, but only clozapine and haloperidol, which reversed amphetamine’s effects on D1-SPN activity, also normalized PPI (**Fig. 3g, h**).

### Chemogenetic D1-SPN inhibition normalizes amphetamine-induced locomotion and PPI deficits

Given that clozapine and haloperidol normalized D1-SPN hyperactivity but the clinically ineffective drug MP-10 did not, we next asked whether modulating D1-SPNs is sufficient to suppress amphetamine-driven changes in locomotion and PPI (**Fig. 2b** and **Fig. 3g**). To do this, we used viruses to express an inhibitory DREADD (DIO-hM4D(Gi)-mCherry) or a control fluorophore (DIO-mCherry) in the DMS of D1-Cre mice^29^ (**Fig. 4a** and **Extended Data Fig. 4a**). The highly selective and brain penetrant DREADD agonist deschloroclozapine^30^ (DCZ) suppressed current-induced D1-SPN spiking in brain slices from experimental mice (**Extended Data Fig. 4b–d**). DCZ treatment (10 µg·kg^-1^) also attenuated amphetamine-driven hyperlocomotion and deficits in PPI in experimental, but not control mice (**Fig. 4b, c**). These effects were less pronounced than systemic clozapine or haloperidol treatment, which may reflect the fact that our virus injections were only in the DMS and did not transduce every D1-SPN (**Extended Data Fig. 4a**). These results imply that counteracting the effects of a hyperdopaminergic state by modulating the activity of D1-SPNs is sufficient to modulate these behaviors in a manner that is consistent with antipsychotic effect.

**Fig. 4:**
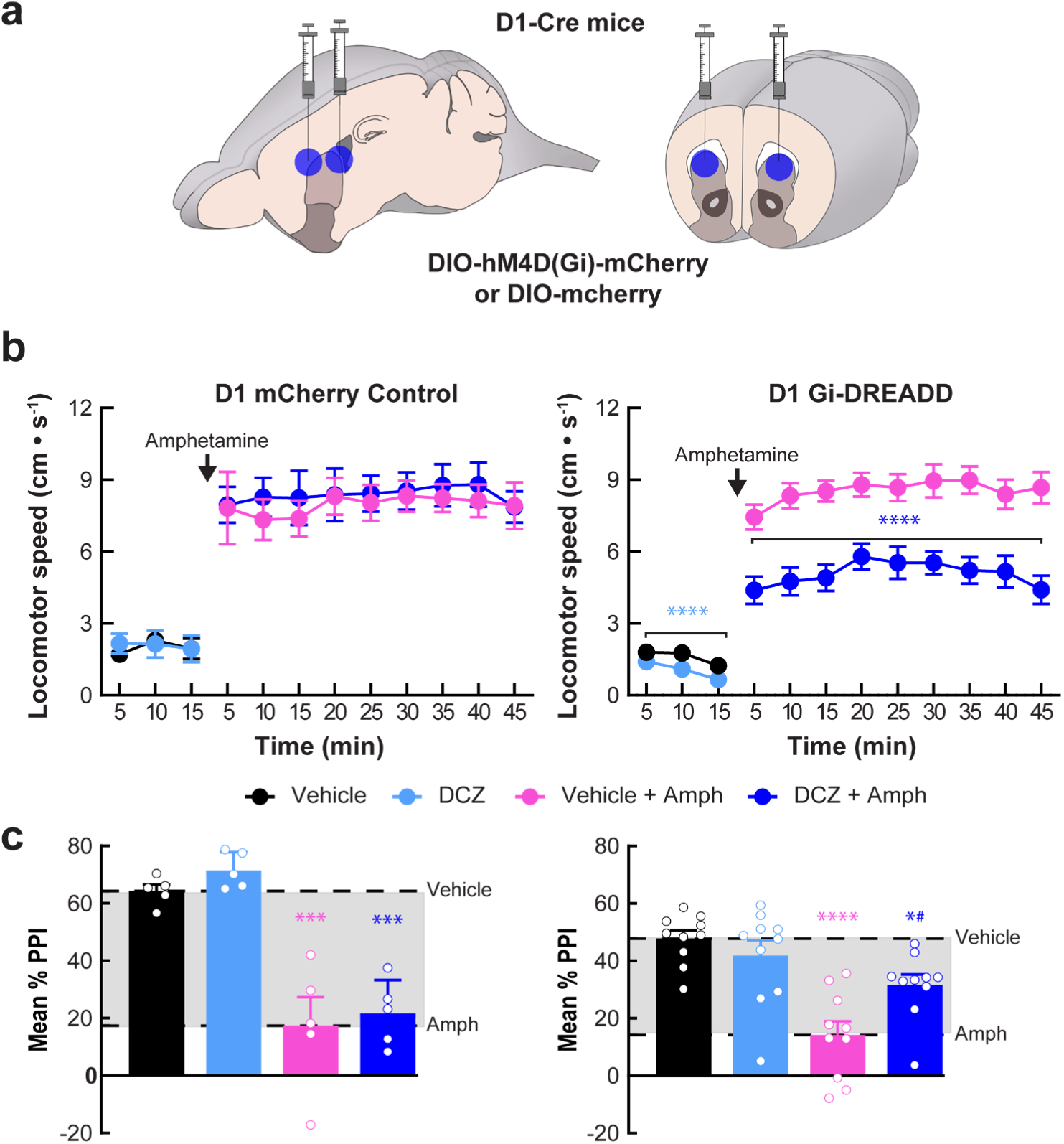
Inhibiting D1-SPNs is sufficient to rescue amphetamine-induced hyperlocomotion and PPI deficits. **a**, We injected DIO-hM4D(G_i_)-mCherry or DIO-mCherry virus bilaterally at two sites in the dorsomedial striatum of D1-Cre mice. **b**, **c**, Treatment with the selective DREADD agonist deschloroclozapine (DCZ) reduced baseline locomotion and attenuated amphetamine-driven hyperlocomotion (**b**) and PPI disruption (**c**) in D1-Cre mice expressing DIO-hM4D(G_i_)-mCherry (*right*), but not the control DIO-mCherry virus (*left*). All data are expressed as mean ± s.e.m. (*N* = 10 experimental and *N* = 5 control mice; \*\*\*\**P* < 0.0001, \*\*\**P* < 0.001, \**P* < 0.05 comparing vehicle or DCZ + amphetamine to vehicle only treatment; ^#^*P* < 0.05 comparing DCZ + amphetamine to vehicle + amphetamine; Holm-Sidak’s multiple comparison test).

### Therapeutically targeting D1-SPNs normalizes their dynamics and behavior

Given that the selective modulation of hyperdopaminergic D1-SPN dynamics was associated with optimal antipsychotic effect, we investigated other therapeutic strategies for targeting these dynamics. Specifically, we focused on three drugs that we predicted would decrease D1-SPN activity under hyperdopaminergic conditions. (1) VU0467154 is a positive allosteric modulator of inhibitory Gαi-coupled M4 muscarinic acetylcholine receptors (M4-PAM) which are specifically expressed in D1-, but not D2-SPNs^24, 31^, (2) SCH23390 is a selective antagonist of excitatory Gαs-coupled D1Rs, and (3) SKF38393 is a D1R partial agonist that we predicted would selectively suppress D1-SPN activity under hyperdopaminergic conditions. We used the same imaging and drug administration procedures to determine how each of these drugs affects D1- and D2-SPN activity under normal and hyperdopaminergic conditions (**Fig. 5a**).

**Fig. 5:**
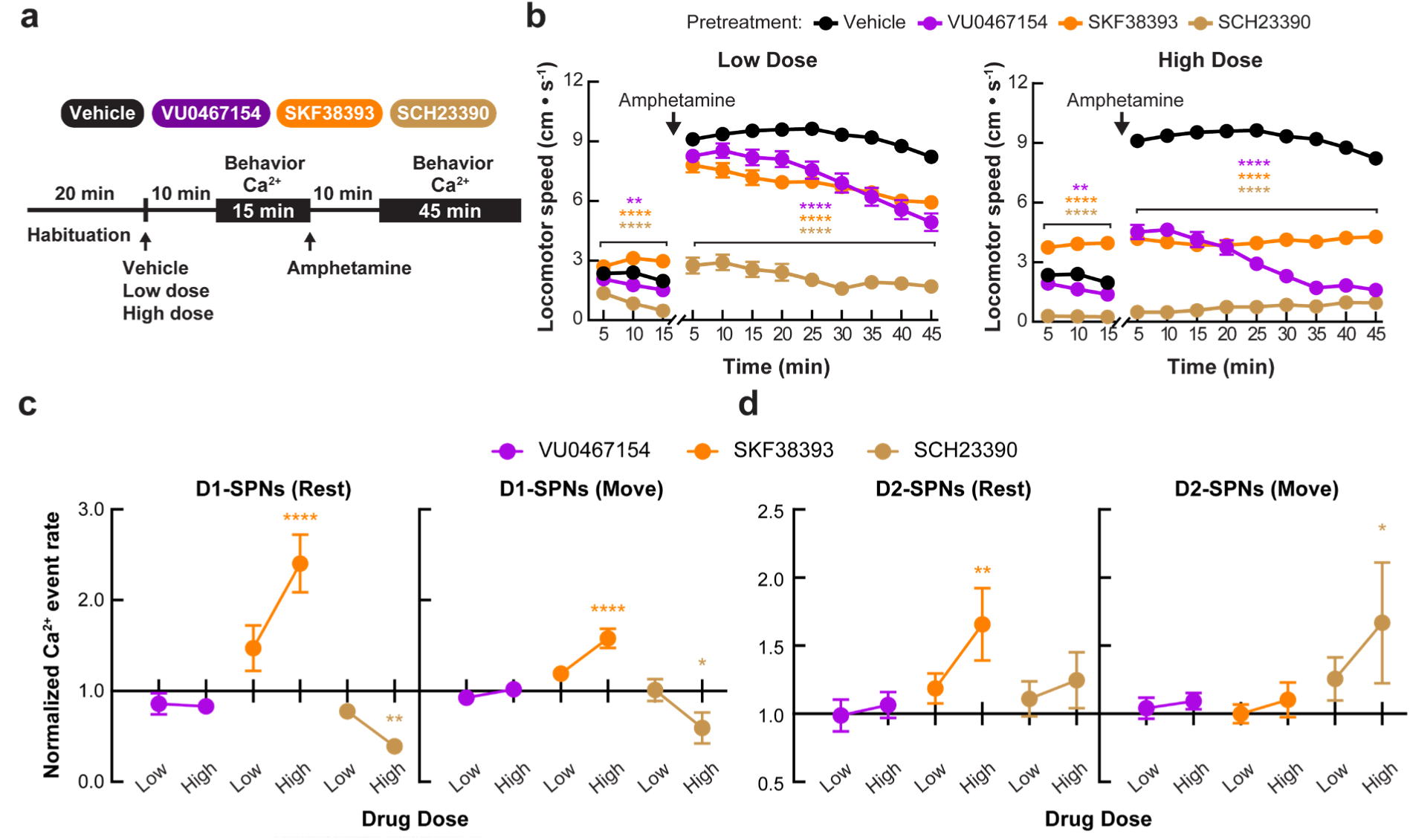
Effects of D1-SPN-targeted compounds on normal D1- and D2-SPN activity levels and spontaneous or amphetamine-driven locomotion. **a**, To record behavior and Ca^2+^ activity, we habituated the mice to the open field arena for 20 min before drug injection. After administering vehicle or a dose of a D1-SPN-targeted drug, we recorded behavior and Ca^2+^ activity for 15 min, administered amphetamine, and recorded Ca^2+^ activity for an additional 45 min. All recordings began 10 min after vehicle, drug, or amphetamine treatment. We administered different drug doses on consecutive days and gave the mice a day off between the different drugs. **b**, Mean ± s.e.m. locomotor speed during the first 15 min recording period following vehicle or drug treatment and the 45 min recording period after amphetamine treatment (*N* = 31 mice; \*\*\*\**P* < 0.0001 and \*\**P* < 0.01 comparing drug to vehicle and drug + amphetamine to vehicle + amphetamine treatment; Holm-Sidak’s multiple comparison test). **c**, **d**, Mean ± s.e.m. Ca^2+^ event rates in D1-SPNs (**c**) and D2-SPNs (**d**) after treatment with a low or high dose of D1-SPN-targeted compounds during rest (*left*) and movement (*right*), normalized to event rates following vehicle only treatment (*N* = 11 D1-Cre and *N* = 10 for A2A-Cre mice; \*\*\*\**P* < 0.0001, \*\**P* < 0.01 and \**P* < 0.05 comparing drug to vehicle treatment; Holm-Sidak’s multiple comparison test).

Under normal conditions, both doses of VU0467154 and SCH23390 decreased, while SKF38393 increased locomotor speed (**Fig. 5b** and **Extended Data Fig. 2b**). Despite reducing locomotion, VU0467154 treatment had no effect on the rates of D1- or D2-SPN activity during either rest or movement. By contrast, the high dose of SCH23390 decreased D1- and increased D2-SPN activity, and the higher SKF38393 dose increased activity levels in both SPN types (**Fig. 5c, d**). Despite the different profiles of their effects on SPN activity levels and locomotion, none of the drugs affected the degree of proximal D1- or D2-SPN activity (**Extended Data Fig. 3c, d**).

Under hyperdopaminergic conditions, pre-treatment with all three drugs dose-dependently reduced hyperlocomotion (**Fig. 5b** and **Extended Data Fig. 2b**). Likewise, all three compounds normalized D1-SPN hyperactivity following amphetamine treatment, though only VU0467154 and SCH23390 also normalized the degree of proximal D1-SPN co-activity (**Fig. 6a, b**). The low dose of SCH23390 was the only treatment that normalized D2-SPN hypoactivity, and none of the compounds had any effects on the spatiotemporal coordination of activity in D2-SPNs (**Fig. 6c, d**). In summary, the three compounds had similar effects on amphetamine-driven hyperlocomotion and the levels of D1-SPN activity, but varied in their profile of effects on other D1- and D2-SPN ensemble dynamics (**Fig. 6e**).

**Fig. 6:**
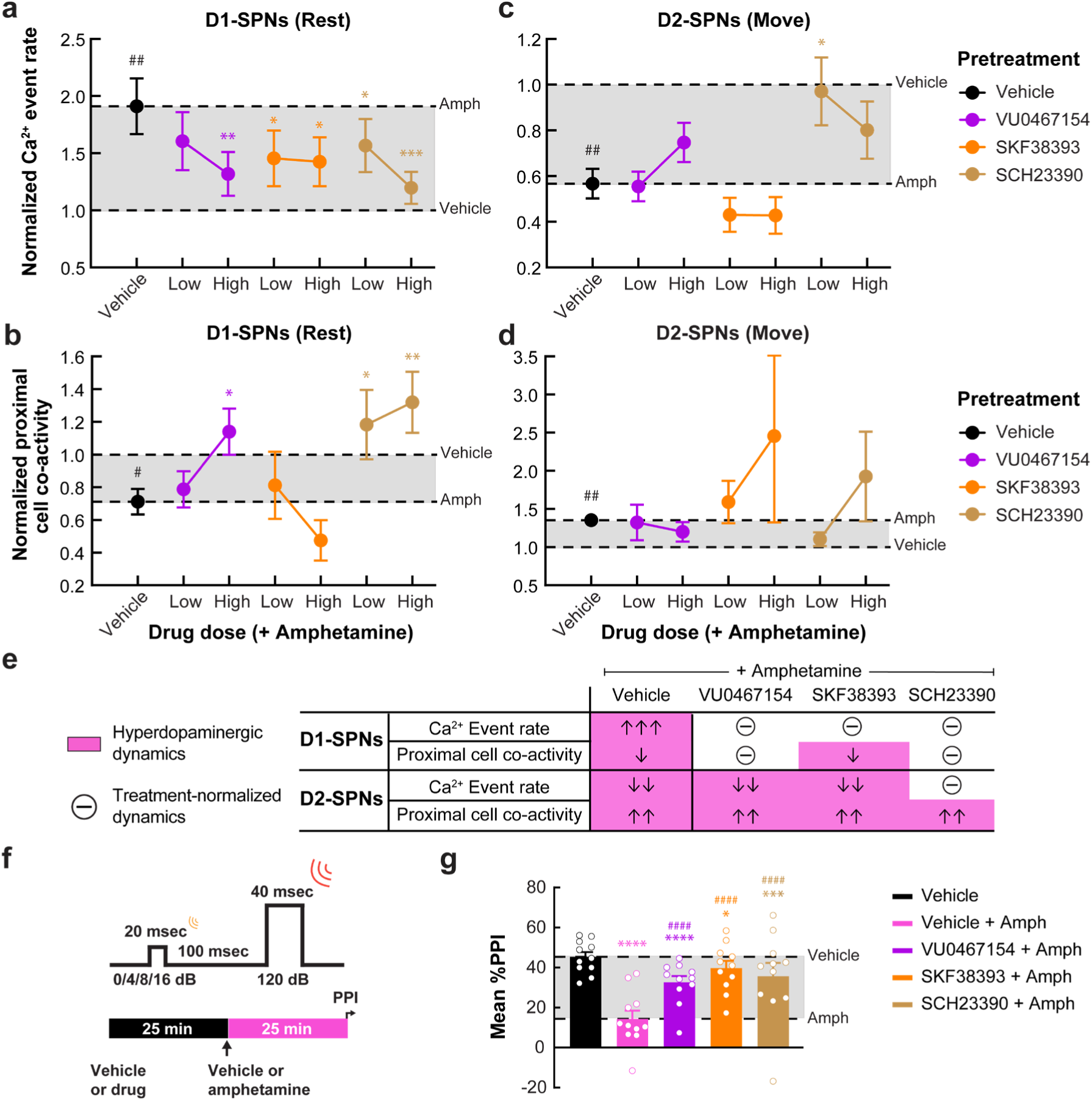
D1-SPN-targeted targeting normalized hyperdopaminergic D1-SPN dynamics and deficits in sensorimotor gating. **a**, **b**, Ca^2+^ event rates (**a**) and proximal co-activity (**b**) of D1- SPNs during periods of rest following vehicle or drug + amphetamine treatment, normalized to values following vehicle only treatment. **c**, **d**, Ca^2+^ event rates (**c**) and proximal co-activity of D2-SPNs (**d**) during periods of movement following vehicle or drug + amphetamine treatment, normalized to values following vehicle only treatment. Data are represented as mean ± s.e.m. (*N* = 11 D1-Cre and *N* = 10 A2A-Cre mice; ^#^*P* < 0.05 and ^##^*P* < 0.01 comparing amphetamine to vehicle treatment; \*\*\**P* < 0.001, \*\**P* < 0.01 and \**P* < 0.05 comparing drug + amphetamine to ve-hicle + amphetamine treatment; Holm-Sidak’s multiple comparison test). **e**, Summary of the effects of different D1-SPN-targeted compounds amphetamine-disrupted D1- and D2-SPN ensemble dynamics. Pink shading indicates neural ensemble dynamics under hyperdopaminergic states. Effect sizes are represented as in Fig. 3e. f, We injected vehicle or D1-SPN-targeted drug 25 min before amphetamine treatment and started measuring PPI 25 min after amphetamine treatment. **g**, Mean ± s.e.m. percent PPI across all pre-pulse intensities following vehicle or drug + amphetamine treatment. (*N* = 11; \*\*\*\**P* < 0.0001, \*\*\**P* < 0.001 and \**P* < 0.05 compared to vehicle treatment; ^####^*P* < 0.0001 compared to vehicle + amphetamine treatment; Holm-Sidak’s multiple comparison test).

Given that D1-SPN suppression was associated with the normalization of PPI under hyperdopaminergic conditions (**Fig. 4c**), we predicted that all three of the D1-SPN-targeted compounds tested here would normalize amphetamine-driven deficits in PPI. Consistent with this prediction, pretreating mice with VU0467154, SCH23390, or SKF38393 all prevented the disruption of PPI by amphetamine (**Fig. 6f, g**). Taken together, these results highlight the potential utility of targeting D1-SPN hyperactivity for antipsychotic effect and delineate logical strategies for doing so.

## Discussion

For decades we have known that dysfunctional dopamine signaling contributes to psychosis and that the efficacy of antipsychotic drugs depends upon their effects on the dopamine system. However, the lack of appropriate tools has hindered our understanding of how dopamine dysfunction and antipsychotic drug treatment affect the function of neural circuits within the dopamine system. Here we applied advanced Ca^2+^ imaging and analysis approaches to define how antipsychotic drugs affect D1- and D2-SPN dynamics *in vivo*, under normal and hyperdopaminergic conditions. Monitoring neural ensemble activity allowed us to differentiate between antipsychotic drugs, even when their effects on mouse behavior were comparable. Further, this approach allowed us to retrospectively identify the neural correlates of known antipsychotic drug efficacy.

Remarkably we found that, despite the longstanding view that antipsychotic drugs work by normalizing D2R signaling and D2-SPN activity, their clinical efficacy was better explained by their ability to normalize D1-SPN activity. This observation led us to explore the therapeutic potential of directly targeting abnormal D1-SPN dynamics. Chemogenetic and directed pharmacological experiments combined with Ca^2+^ imaging confirmed this potential, which has largely been overlooked as a target for therapeutic development. In addition, our imaging results produced other unexpected findings that challenge our understanding of how specific dopamine receptors modulate intact striatal circuit function. These additional findings also unveiled differences between each compound with implications for therapeutic development that warrant further consideration.

### Hyperdopaminergic striatal ensemble dynamics

Our study provides the first detailed analysis of how amphetamine alters the neural ensemble dynamics of D1- and D2-SPNs. Amphetamine treatment predictably enhanced D1- and suppressed D2-SPN activity levels, consistent with earlier studies reporting heterogeneous effects of amphetamine on unidentified SPN activity levels^32^. Unexpectedly these effects were dependent on locomotor state. These state-dependent effects may coincide with earlier reports of dose-dependent amphetamine treatment effects on SPN activity, whereby lower doses increase and higher doses suppress SPN activity levels^33^. Given the correlations between amphetamine treatment dose, dopamine transmission levels and locomotor speed, the selective increase in D1-SPN activity at lower speeds and decrease in D2-SPN activity at higher speeds may reflect differences in striatal dopamine levels at different locomotor states (**Fig. 1c–e**).

Amphetamine treatment also differentially altered the degree of spatially coordinated D1- and D2-SPN co-activity (**Fig. 1f–h**). We previously observed de-correlated D1-SPN hyperactivity during L-DOPA-induced dyskinesia, which is intriguing given that dyskinesia and disorganized behavior also occur in schizophrenia, even in drug-naïve patients^12, 34^. These observations suggest that the neural substrates of psychosis and dyskinesia may both result from excess dopaminergic modulation of intrinsic excitability or synaptic strength in D1-SPNs. By contrast, D2-SPNs increased their spatiotemporal coordination following amphetamine treatment. This was a novel signature of striatal network dysfunction that was not observed in our earlier studies^12^. This heightened co-activation may reflect the diminution of lateral inhibition between D2- SPN cell pairs, possibly via axon-terminal D2R activation^35^. While this neural ensemble signature was unique to this study, the fact that neither haloperidol nor clozapine altered proximal D2- SPN co-activity following amphetamine treatment argues against a causal role of these dynamics in psychosis.

### The neural ensemble correlates of antipsychotic drug efficacy

We tested two efficacious antipsychotic drugs (clozapine and haloperidol) and one inefficacious drug candidate (MP-10). Under hyperdopaminergic conditions, clozapine only affected D1-SPNs, haloperidol affected both SPN types, and MP-10 normalized D2- but exacerbated hyperdopaminergic D1-SPN dynamics (**Fig. 3e**). Clozapine’s superior clinical efficacy and favorable side-effect profile have long been recognized, but incompletely understood. Clozapine is the prototype of the atypical antipsychotic drugs, which are distinguished by their lower D2 and higher 5-HT2 receptor family affinities^36^. This pharmacological profile is thought to permit antipsychotic effect with a lower level of D2R engagement, precluding the adverse (*e.g.*, motor) effects associated with first-generation antipsychotics like haloperidol^37, 38^. In addition, this unique pharmacology is thought to underlie clozapine’s superior efficacy for psychosis, including its treatment-resistant manifestations^39, 40^. This complex pharmacology, including the fact that clozapine binds to D1Rs *in vivo*^41^, made it difficult to predict but less of a surprise that clozapine only affected hyperdopaminergic D1-SPN dynamics. Likewise, although haloperidol’s greater specificity for D2Rs explains its effects on D2-SPN activity, its normalization of hyperdopaminergic D1-SPN dynamics was also surprising. However, D2 and other brain receptors bound by haloperidol are located throughout the cortico-basal ganglia-thalamic circuit, including in local striatal interneurons, which could indirectly contribute to haloperidol’s effects on D1-SPN activity^42-50^.

Perhaps even more surprising than clozapine and haloperidol’s effects on D1-SPN activity was the fact that the clinically non-efficacious drug MP-10 completely normalized hyperdopaminergic D2-SPN dynamics (**Fig. 3c**, **d**). At face value, this finding implies that normalizing D2-SPN activity is not sufficient to produce an antipsychotic effect. However, both D1- and D2-SPNs express PDE10A, the enzyme inhibited by MP-10^51^. Still, previous reports suggest that PDE10A inhibition preferentially affects D2-SPNs^52, 53^. Consistent with this idea, MP-10 increased D2-, but not D1-SPN activity levels under normal conditions (**Fig. 2c, d**). However, MP-10 treatment exacerbated the D1-SPN hyperactivity observed under hyperdopaminergic conditions (**Fig. 3a**). Therefore, one possible explanation for MP-10’s lack of antipsychotic effects in patients is that its effects on D1-SPN activity counteracts any of its therapeutic effects on D2-SPN dynamics. Future experiments determining whether other effective antipsychotics exclusively act on D2-SPNs under hyperdopaminergic conditions will be necessary to demonstrate whether solely modulating D2-SPNs is associated with clinical therapeutic effect of D2-SPN modulation. Nevertheless, our current results suggest that normalizing D1-SPN dynamics is sufficient for antipsychotic effect, and that selectively doing so may be optimal.

### Therapeutically targeting hyperdopaminergic D1-SPN dynamics

Given the apparent link between normalizing D1-SPN dynamics and antipsychotic effect, we next asked whether we could selectively target these dynamics therapeutically. Chemogenetically inhibiting D1-SPNs in the DMS was sufficient to normalize amphetamine-driven changes in locomotion and sensorimotor gating. Likewise, three D1-SPN-targeted compounds all normalized D1-SPN hyperactivity, locomotion, and sensorimotor gating following amphetamine treatment. The first of these compounds was VU0467154, positive allosteric modulator (PAM) of Gi/o-coupled M4 cholinergic receptors, which are expressed in D1-, but not D2-SPNs^31^.

VU0467154 or other M4-PAMs are known to have antipsychotic-like effects on behavior in animal models related to psychosis^24, 54, 55^. In our neural ensemble readout of drug efficacy, VU0467154 was the most similar to clozapine, in that it selectively normalized de-correlated D1-SPN hyperactivity following amphetamine treatment, but had no effect on D2-SPNs (**Fig. 6e**). The second drug we tested, the D1R partial agonist SKF38393, also only affected hyperdopaminergic D1-SPN activity levels, but in contrast to VU0467154 it failed to normalize the spatiotemporal dynamics of D1-SPN activity. By contrast, the D1R antagonist SCH23390 was most similar to haloperidol, in that it normalized the hyperdopaminergic dynamics of both D1- and D2-SPNs.

Over the past decade, much of the focus on D1Rs in schizophrenia has been on augmenting their signaling to promote cognition^56–59^. This idea is largely based on foundational work showing that, in schizophrenia, dopamine transmission is decreased within the prefrontal cortex, where D1Rs are enriched and their signaling is crucial for cognitive function^60, 61^. This results in regional imbalance in dopamine signaling between cortex and striatum that poses a challenge to therapeutic development for schizophrenia. For example, attenuating dopamine signaling may be crucial for treating psychosis, but treatments that do so may exacerbate the cognitive symptoms of schizophrenia. Likewise, augmenting dopamine signaling might improve cognition, but our results suggest that doing so, particularly at D1Rs, could also worsen the cardinal symptoms of psychosis.

Dopamine stabilizers, such as the partial D2R agonist aripiprazole, are considered to be a possible solution to this problem, by exerting state-dependent effects on dopamine receptor signaling^62, 63^. Specifically, aripiprazole is thought to act as a D2R agonist under conditions of low dopamine (*i.e.*, in cortex) and antagonist under high dopamine conditions (*i.e.*, striatum). Given the importance of cortical D1R signaling for cognition and the association between normal D1-SPN dynamics and antipsychotic effect uncovered here, we reasoned that a D1R partial agonist might better stabilize dopamine signaling across cortex and striatum and in different dopaminergic states. Consistent with this idea, SKF38393 exhibited dopamine agonist-like effects on D1-SPN activity under normal conditions, but suppressed D1-SPN activity following amphetamine treatment, similar to the D1R antagonist SCH23390 (**Fig. 5c**; **Fig. 6a**).

SCH23390 normalized more of the amphetamine-driven changes in D1- and D2-SPN ensemble dynamics than the other two D1-SPN-targeted drugs, but exerted clear D1R antagonist effects on D1-SPN activity levels under normal conditions (**Fig. 5c**; **Fig. 6a–c**). Given that an antagonist is predicted to block D1R signaling at both low/normal as well as high dopamine conditions, D1R antagonism may have limited utility for treating regional dopamine dysfunction in schizophrenia. Consistent with this idea, D1R antagonism does not appear to be an effective therapeutic strategy for psychosis^64^. By contrast, modulating M4Rs provides a possible approach to stabilize D1R signal transduction in the striatum without specifically counteracting D1R signaling under normal or low dopamine conditions. Consistent with this idea, VU0467154 had no effect on D1- or D2-SPN under normal conditions, but completely normalized D1-SPN dynamics following amphetamine treatment (**Fig. 5c, d; Fig. 6a, b**). Taken together, our results suggest that M4 positive allosteric modulation and possibly D1R partial agonism might provide similar therapeutic effects to clozapine, with the minimal motor side effects and the potential for the regional stabilization of brain dopamine function. Intriguingly, D1R agonism is predicted to improve cognitive function under hypodopaminergic conditions in the cortex, and an M4-PAM has been shown to have pro-cognitive effects in animal models related to schizophrenia^54^.

### Neural ensemble correlates of adverse drug effects

Although we primarily considered efficacy in terms of each drug’s effects under hyperdopaminergic conditions, each drug’s effects on D1- and D2-SPN ensemble dynamics under normal conditions is another important therapeutic consideration. In schizophrenia patients, fluctuations in striatal dopamine are thought to drive psychotic episodes, and dopamine transmission is normal in patients with stable symptoms^65^. Taking this into consideration, the ideal treatment would minimally affect striatal activity under normal conditions, but block the effects of excess dopamine transmission during psychotic episodes. Of particular relevance to antipsychotic drugs is their propensity for adverse motor effects, such as parkinsonism. We recently used this approach to characterize the D1- and D2-SPN ensemble correlates of parkinsonism following the chemical lesion of dopamine neurons^12^. Specifically, we found that the loss of dopamine decreases D1- and increases D2-SPN activity levels. Among the drugs tested here, clozapine and VU0467154 were the only ones that did not increase D2-SPN activity levels under normal conditions.

SCH23390 additionally reduced D1-SPN activity under normal conditions, suggesting D1R antagonism may have a particularly high propensity for adverse motor effects (**Fig. 3d**; **Fig. 5c, d**). These parkinsonism-associated dynamics may underlie the different motor side-effect propensities of drugs like haloperidol and clozapine, and help predict the side-effect propensities of other candidate treatments.

Overall, the neural ensemble approach described here is a powerful tool for understanding the mechanisms underlying brain diseases and their effective treatment. We demonstrated the utility of this approach for predicting the different efficacies and side-effect propensities of three antipsychotic drugs. Our results suggest that the optimal therapy for psychosis specifically normalizes D1-SPN dynamics under hyperdopaminergic conditions and minimally perturbs D1- and D2-SPN activity under normal conditions. We found that normalizing D1-SPN hyperactivity is sufficient to rescue amphetamine-driven disruption of antipsychotic responsive behaviors, and we adjudicate three potential therapeutic strategies for targeting aberrant D1-SPN dynamics. These findings have the potential to inform the development of novel treatments for psychosis with fewer adverse effects and greater overall efficacy.

## Methods

### Mice

All mice were housed and handled according to guidelines approved by the Northwestern University Animal Care and Use Committee. We used both male and female mice for all experiments. For Ca^2+^ imaging and DREADD experiments, we used GENSAT *Drd1a* (FK150) or *Adora2a* (KG139) BAC transgenic Cre-driver mouse lines from the Mutant Mouse Research & Resource Centers (www.mmrrc.org), backcrossed to a C57BL/6J background (Jax # 000664). For PPI experiments with amphetamine + antipsychotic drug treatment, we used C57BL/6J mice. All mice were 12–24 weeks at the start of experimental testing, with the exception of the mice used for slice electrophysiology, which were aged 7–8 weeks at the time of testing.

### Virus injections

For all surgeries, we anesthetized mice with isoflurane (2% in O_2_) and stereotaxically injected virus at a rate of 250 nL·min^-1^ into the DMS using a microsyringe with a 33-gauge beveled tip needle. All anterior-posterior (AP) and medial-lateral (ML) coordinates are reported from bregma, while all dorsal-ventral (DV) coordinates are reported from dura. For all DV coordinates, we went 0.5-mm past the injection target, and then withdrew the syringe back to the target for the injection. After each injection, we left the syringe in place for five min, withdrew the syringe 0.1 mm and waited five more min before slowly withdrawing the syringe completely. Following virus injection we sutured the scalp, injected analgesic (Buprenorphine SR; 1 mg·kg^-1^), and allowed the mice to recover for a week before implanting an optical guide tube.

For Ca^2+^ imaging experiments, we injected 500 nL of AAV2/9-Syn-FLEX-GCaMP7f (1.6 × 10^12^ GC·mL^-1^; AP: 0.8 mm, ML: 1.5 mm and DV: -2.7 mm). To transduce a wider range DMS neurons for DREADD behavioral experiments, we injected 650 nL of AAV2/9-hSyn-DIO-hM4Di-mCherry (5.0 × 10^12^ GC·mL^-1^) or AAV2/9-hSyn-DIO-mCherry (1.15 × 10^12^ GC·mL^-1^) bilaterally at two sites in each hemisphere (AP: 0.4 mm, ML: ±1.5 mm and AP: 1.2 mm, ML: ±1.25, both DV: -2.8 mm). For sparser transduction in our DREADD electrophysiology experiments, we injected 650 nL of AAV2/9-hSyn-DIO-hM4Di-mCherry (1.25 × 10^12^ GC·mL^-1^) bilaterally at two sites in each hemisphere (AP: 0.4 mm, ML: ±1.5 mm and AP: 1.2 mm, ML: ±1.1 mm, both DV: -2.5 mm). We obtained all viruses from AddGene.

### Implant surgeries

We constructed optical guide tubes by using ultraviolet (UV) liquid adhesive (Norland #81) and a UV spot curing system (Electro-Lite) to fix a 2-mm-diameter disc of #0 glass (TLC International) to the tip of a 3.8-mm-long, 18-gauge, extra-thin stainless steel tube (Ziggy’s Tubes and Wires). We ground off any excess glass using a polishing wheel (Ultratec).

To prepare mice for Ca^2+^ imaging, we anesthetized virus-injected mice with isoflurane (2% in O2) and a 1.4-mm-diameter drill bit was used to create a craniotomy (AP: 0.8 mm; ML: 1.5 mm) for implantation of the optical guide tube. We used a 0.5-mm diameter drill bit to drill four additional small holes at spatially distributed locations for insertion of four anchoring skull screws (Antrin miniature specialties). We aspirated cortex down to DV: -2.1 mm from dura by using a 27-gauge blunt-end needle and implanted the optical guide tube at DV: -2.35 mm from dura. After placing the guide tube, we applied Metabond (C&B Metabond) to the skull, and then used dental acrylic (Coltene) to fix the full assembly along with a stainless steel head-plate (Laser Alliance) for head-fixing mice during attachment and release of the miniature microscope. We injected analgesic (Buprenorphine SR; 1 mg·kg^-1^) and allowed the mice to recover for 3–4 weeks before mounting the miniature microscope.

### Miniature microscope mounting

We head-fixed each guide-tube implanted mouse by its headplate on a running wheel and inserted a gradient refractive index (GRIN) lens (1-mm diameter; 4.12-mm length; 0.46 numerical aperture; 0.45 pitch; Inscopix Inc.) into the optical guide tube. We then assessed GCaMP7f expression in the DMS using a commercial two-photon fluorescence microscope (Ultima Investigator, Bruker). We then anesthetized mice with substantial GCaMP7f expression with isoflurane (2% in O_2_), placed them back into the stereotaxic frame, and glued the GRIN lens in the guide tube with UV light curable epoxy (Loctite 4305). Next, we used the stereotaxic manipulator to lower the miniature microscope with its attached base plate (nVistaHD, Inscopix Inc.) toward the GRIN lens until the fluorescent tissue came into focus. We then created a structure of blue-light curable resin (Flow-It ALC; Pentron) on the dental acrylic skull cap around the base plate, then attached the structure to the miniature microscope base plate using UV curable epoxy (Loctite 4305). Finally, we coated the epoxy/resin with black nail polish to make it opaque.

### *In vivo* pharmacology

We administered all drugs via subcutaneous injection (10 mL·kg^-1^ injection volume) on sequential days at the escalating dosages and order depicted in **Fig. 2a** and **Fig. 5a**. All mice received one day off between treatments with the different drugs. We dissolved clozapine (2 or 3.2 mg·kg^-1^) and haloperidol (0.1 or 0.32 mg·kg^-1^) in 0.3% tartaric acid. We dissolved SCH23390 (0.032 or 0.1 mg·kg^-1^) and D-Amphetamine hemisulfate (2.5 or 10 mg·kg^-1^) in saline (0.9% NaCl). We dissolved MP-10 (1 or 3.2 mg·kg^-1^) in 5% 2-Hydroxypropyl-β-cyclodextrin in saline, VU0467154 (1 or 10 mg·kg^-1^) in 10% Tween 80, SKF38393 (10 or 100 mg·kg^-1^) in water, and DCZ (10 µg·kg^-1^) in 2% DMSO. We obtained VU0467154 from the Vanderbilt Center for Neuroscience and Drug Discovery, DCZ from MedChemExpress, and all other drugs from Sigma.

To examine the effects of vehicle or antipsychotic drugs under normal and hyperdopaminergic states, we injected each drug or its corresponding vehicle and waited 10 min before recording open field behavior + Ca^2+^ activity for 15 min. We then injected amphetamine (2.5 mg·kg^-1^) and waited 10 min before recording behavior + Ca^2+^ activity for 45 min (**Fig. 2a** and **Fig. 5a**). For PPI experiments, we administered the higher of the two doses of each drug or their corresponding vehicle 25 min before amphetamine injection (10 mg·kg^-1^) and measured PPI 25 min after amphetamine treatment (**Fig. 3f** and **Fig. 6f**). For chemogenetic experiments in the open field, we administered DCZ or its vehicle 10 min before recording behavior for 15 min, then administered amphetamine (2.5 mg·kg^-1^) and waited 10 min before recording behavior for 45 min. For chemogenetic experiments during PPI, we administered DCZ 25 min before amphetamine injection (10 mg·kg^-1^) and measured PPI 25 min after amphetamine treatment.

### *In vivo* Ca^2+^ imaging

We habituated mice to a circular open field arena (30.48-cm diameter) for three days (1 h per day), during which time we also habituated the mice to two subcutaneous injections of saline and one injection of amphetamine (2.5 mg·kg^-1^). Just before each Ca^2+^ imaging session, we briefly head fixed the mouse by its implanted head plate on a running wheel. We then attached the miniature microscope, adjusted its focal plane, and then released the mouse after securing the microscope. After 20 min habituation in the open field, we injected mice with vehicle or drug, waited 10 min and recorded Ca^2+^ activity for 15 min, then injected amphetamine, waited 10 min, and recorded Ca^2+^ activity for 45 min as described in **Pharmacology (Fig. 2a** and **Fig. 5a)**. We used an illumination power of 50–200 µW at the specimen plane and a 20-Hz image frame-acquisition rate.

#### PPI

We placed mice into a plexiglass cylinder (10 × 20 × 10 cm) on a platform equipped with a piezoelectric transducer inside of a larger, sound-attenuating chamber with 65 dB of continuous background noise (SR-Lab; San Diego Instruments). Mice received 2 × 30 min habituation sessions on two consecutive days. During experimental testing, we treated mice with vehicle, drug, or DCZ + amphetamine (as described in **Pharmacology**) and placed them into the startle chamber. Evaluation of PPI consisted of 5 min acclimation followed by five priming acoustic stimulus pulses (120 dB; 40 ms) then 20 trial blocks of pseudo-randomly presented trials of no-stimulus pulse or pre-pulse (0, 4, 8, or 16 dB above background; 20 ms) 100 ms before the acoustic startle stimulus (120 dB; 40 ms) (**Fig. 3f** and **Fig. 6f**). The intertrial interval (ITI) averaged 17 s (range 10–25 s). We calculated the levels of PPI at each pre-pulse intensity as 100 - [100 × (response amplitude for each pre-pulse stimulus with startle stimulus) / (response amplitude for 0 dB pre-pulse with startle stimulus)]. Mean % PPI was calculated by averaging levels of PPI at each pre-pulse intensity level.

### Histology

After all behavioral, Ca^2+^ imaging, and PPI experiments, we euthanized and intracardially perfused the mice with phosphate buffered saline (PBS) and then a 4% solution of paraformaldehyde in PBS. We sliced 80-µM-thick coronal sections from the fixed-brain tissue using a vibratome (Leica VT1000s). For immunostaining, we used an anti-GFP antibody (1:1000, Invitrogen, A11122) and a fluorophore-conjugated secondary antibody (1:500, Jackson Immunoresearch 711-546-152), then mounted the sections with DAPI-containing fluoromount (SouthernBiotech, 0100-20). We imaged slices using a fluorescent microscope (Keyence BZ-X800) with a 10x objective.

### Slice electrophysiology

We anesthetized and transcardially perfused mice with ice-cold, carbogen-saturated cutting solution (185 mM sucrose, 2.5 mM KCl, 25 mM NaHCO_3_, 1.25 mM NaH_2_PO_4_, 0.5 mM CaCl_2_, 10 mM MgCl_2_, and 25 mM glucose, pH 7.3 [315–320 mOsm·L^-1^]). Following perfusion, we decapitated the mice, rapidly removed the brain and sectioned it in an ice-cold carbogen-saturated cutting solution using a vibratome (VT1000S, Leica Microsystems). We then incubated coronal slices (220 µm) in carbogen-saturated artificial cerebrospinal fluid (ACSF) containing 93 mM NMDG, 93 mM HCl, 2.5 mM KCl, 30 mM NaHCO_3_, 1.2 mM NaH_2_PO_4_, 20 mM HEPES, 5 mM Na-ascorbate, 3 mM Na-pyruvate, 2 mM thiourea, 0.5 mM CaCl_2_, 10 mM MgSO_4_, and 25 mM glucose, pH 7.3 (315–320 mOsm**·**L^-1^) at 32–34°C for 10 min, then in carbogen-saturated ACSF containing 125 mM NaCl, 2.5 mM KCl, 25 mM NaHCO_3_, 1.25 mM NaH2PO_4_, 2 mM CaCl_2_, 1 mM MgCl_2_, and 25 mM glucose, pH 7.3 (315–320 mOsm L^-1^) at room temperature for at least 1 hr before electrophysiological recordings. We transferred the brain slices to a small-volume (< 0.5 ml) recording chamber mounted on a fixed-stage, upright microscope. We performed all electrophysiological recordings at 32–34°C. The chamber was superfused with carbogen-saturated ACSF (SH-27B with TC-324B controller, Warner Instruments). We performed conventional whole-cell patch-clamp recordings on visually identified (60 X, 0.9 NA water-immersion objective) D1-SPNs expressing mCherry. Recording electrodes had tip resistances of 3-8 MΩ when filled with internal recording solution containing (in mM): 125 KMeSO4, 5 KCl, 5 NaCl, 0.02 EGTA, 11 HEPES, 1 MgCl, 10 phosphocreatine-Na_2_, 4 Mg-ATP, 0.3 Na-GTP, adjusted to pH 7.2, 300 mOsm·L^-1^. We made all recordings using MultiClamp 700B amplifiers and filtered all signals at 2 kHz and digitized at 10 kHz. We discarded data if the input resistance changed >20% over the time course of the experiment. For drug treatment, we perfused vehicle (0.2% DMSO), DCZ (100 nM or 1 µM) or 10 µM of clozapine-N-oxide (CNO) for 1 mL·min^-1^.

### Behavioral tracking

We used a TTL-triggered video camera with IC Capture 2.4 software (The Imaging Source) with a varifocal lens (T3Z2910CS; Computar) to record 20-Hz videos of freely moving mouse behavior. We used software written in ImageJ and part of the CIAtah analysis suite (https://ba-hanonu.github.io/ciatah/) to track each mouse’s position in an open field arena. Briefly, we used this software to identify the mean location of the largest, and darkest contiguous pixel group (*i.e.*, the mouse) in each movie frame, then computed the mouse’s locomotor speed from the trajectory of its centroid location across movie frames. We then applied a 1-s median filter to the resulting speed trace and down-sampled the trace by a factor of 4 to match the temporal resolution of our 5-Hz Ca^2+^ recordings. We classified each 5-Hz time bin of the speed trace as one in which the mouse was either ‘resting’ or ‘moving’, according to whether its instantaneous speed was below or above 0.5 cm·s^-1^, respectively. If the mouse was ‘moving’ in two time bins separated by <1 s, we classified the intervening time bins as ones in which the mouse was ‘moving’.

### Ca^2+^ movie pre-processing

We used the CIAtah analysis suite to 1) down-sample the acquired Ca^2+^ movies in space using 2 × 2 bi-linear interpolation, 2) reduce background fluorescence by applying a Gaussian low-pass spatial filter to each movie frame and dividing each frame by its low-pass filtered version, 3) motion correct using the TurboReg algorithm, 4) normalize each movie by subtracting the mean fluorescence value for each pixel in time and dividing each pixel by the same mean fluorescence [(*F*(*t*) - *F*_0_)*/F*_0_], and 5) temporally down-sample the resulting Δ*F/F* movies by a factor of 4 using linear interpolation to a frame rate of 5 Hz.

### Active neuron identification

We used Constrained Nonnegative Matrix Factorization for microendoscopic data (CNMF-E)^66^ to extract putative neurons from the processed Δ*F/F* movies. We then visually inspected and manually classified candidate cells in 12% of the Ca^2+^ imaging sessions (42 of 361 total imaging sessions) based on their size, shape, and Ca^2+^ activity trace. We then used these manually sorted data to train a machine-learning based classifier (using the CLEAN module in CIAtah) for automated sorting of the entire data set. This automated classifier categorized candidate cells based on the evaluation of 21 features of the CNMF-E spatial filters, their Ca^2+^ activity traces, and the Δ*F/F* movies. Parameters included: the 1) diameter 2) area and 3) perimeter of the cellular filter; (4) proportion of the pixels in the convex hull that were also in the spatial filter; the (5) skewness and (6) kurtosis of the statistical distribution of intensity values in the spatial filter; (7) mean value of the signal-to-noise ratio (SNR), averaged over all Ca^2+^ transients within the candidate cell; number of Ca^2+^ transients greater than (8) 1, (9) 3, and (10) 5 times the s.d. of the noise fluctuations within the candidate cells; (11) mean ratio of the peak rise and decay slopes of the Ca^2+^ transients; (12) mean full-width half max value of the Ca^2+^ transients; (13) mean amplitude of the Ca^2+^ transients; the (14) skewness and (15) kurtosis of the statistical distribution of intensity values of the full Ca^2+^ activity trace for each candidate cell; (16) mean amplitude variance at each time point in a 16-s-window around each Ca^2+^ transient waveform; (17) mean correlation coefficient of all Ca^2+^ transient waveforms; (18) the mean correlation coefficient between the CNMF-E image and, at most, 10 images taken from frames temporally aligned to Ca^2+^ event transients in the movie and cropped to a 20 × 20 pixel region centered on the CNMF-E image centroid; (19) the same as (18) but using a binarized image (all pixels below 40% of the maximum value set to zero, all above set to one); (20) the same as (18) but using only the maximum correlation coefficient from all CNMF-E-movie frame image comparisons; and (21) the same as (19) but using only the maximum correlation coefficient from all CNMF-E movie frame image comparisons. After computing these parameters for every candidate cell identified by CNMF-E, we used MATLAB’s *Statistics and Machine Learning* and *Deep Learning* software toolboxes to train support vector machine (SVM), general linear model (GLM), and neural network (nnet) classifiers to automatically classify neurons in our data set.

### Ca^2+^ event detection

After extracting all individual cells and their time traces of Ca^2+^ activity, we evaluated the individual Ca^2+^ events in each cell’s time trace using a threshold-crossing algorithm^67^. Noise and reduced fluctuations in baseline fluorescence were removed by averaging over a 600 ms (3 frame) sliding window, then subtracting a median-filtered version (40 s sliding window) of the trace from the smoothed version. We calculated the standard deviation (s.d.) of the resulting trace and identified any peaks that were ≥2.5 s.d. above baseline noise while enforcing a minimum interevent time of >1.6 s. We determined the initiation time of each Ca^2+^ event as the temporal mid-point between the time of each event’s fluorescence peak and the most recent preceding trough in fluorescence. All subsequent data analyses of neural activity used the resulting 5-Hz binarized event trains in which a ‘1’ indicated the initiation of a Ca^2+^ event. To generate the illustrative Ca^2+^ activity traces in **Fig. 1b**, for each example cell we set to zero all pixels of the cell’s spatial filter with weights <50% of the maximum value in the filter, and then applied the truncated filter to the Δ*F/F* movie to generate a Ca^2+^ activity trace.

### Analysis of pairwise cell co-activity

To identify correlated Ca^2+^ activity within each frame, we evaluated the fraction of all Ca^2+^ events in the two cells in each frame. This fraction is equivalent to a Jaccard index, *J*, of the two cells’ correlated activity (*J* = |T_1_ ∩ T_2_| / |T_1_ ∪ T_2_|), where T_1_ and T_2_ are the binarized rasters of Ca^2+^ events for the two cells^12^. We plotted for all cell pairs the Jaccard index as a function of anatomical separation between the two cells’ centroids. To control for any effects of time-varying Ca^2+^ event rates on the Jaccard indices, we also computed the Jaccard indices for datasets in which the binarized Ca^2+^ event trace for each cell was circularly permuted in time by a randomly chosen temporal displacement. We did this for 1000 different randomly permutated datasets. We then normalized the Jaccard index values in the real data by those obtained from the shuffled datasets. We defined ‘proximal cell co-activity’ as the mean Jaccard index for cell pairs within 25– 125 µm, normalized by the corresponding value of the shuffled datasets (**Extended Data Fig. 1c**). To examine the relationship between proximal cell co-activity and locomotor speed, we sub-divided the shuffle-normalized proximal jaccard indices into bins corresponding to locomotor speeds ranging from 0.5–14 cm·s^-1^. The bin sizes ranged from 0.5 cm·s^-1^ bins at lowest speeds to 6 cm·s^-1^ bins at highest speeds, for which the statistical sampling was sparse (**Fig. 1f**; **Extended Data Fig. 1d**). To compare drug effects to vehicle, we normalized the values in each speed bin to the corresponding values following vehicle or vehicle + amphetamine treatment, then averaged the speed bins during periods of rest and movement (locomotor speeds < 0.5 cm·s^-1^ and >= 0.5 cm·s^-1^, respectively; **Fig. 1h**; **Fig. 3b, d; Fig. 6b, d**; **Extended Data Fig. 3a–d**).

### Analysis of event rates

We used the binarized Ca^2+^ event traces of each cell’s activity to compute each cell’s Ca^2+^ event rate as a function of locomotor speed using the same speed bins described above (**Fig. 1c**; **Extended Data Fig. 1b**). As with the proximal cell co-activity, we normalized the values in each speed bin to the corresponding values following vehicle or vehicle + amphetamine treatment, then averaged the speed bins during periods of rest and movement (**Fig. 1e**; **Fig. 2c, d**; **Fig. 3a, c**; **Fig. 5c, d**; **Fig. 6a, c**).

### Data analysis and statistical tests

We performed data analysis using custom software written in MATLAB and ImageJ. We used Prism (GraphPad) to perform statistical tests. We used two-tailed, non-parametric statistical tests to avoid assumptions of normal distributions and equal variance across groups. For paired tests, we used Wilcoxon signed-rank tests or one-way repeated measures ANOVA. For drug × dose comparisons, we used two-way repeated measures ANOVA. In some cases, individual doses were missing due to errors in the recording on those days. In those cases, we used a mixed effects model to evaluate drug × dose comparisons. For post hoc tests, we used a Holm-Sidak correction for multiple comparisons. *P*- and *N-*values for the statistical tests are provided in **Supplementary Table 1**.

### Data availability

The software code used to process our Ca^2+^ movies are freely available (https://ba-hanonu.github.io/ciatah/). The data and any custom scripts that support our findings are available upon request to the corresponding author.

## Acknowledgements

We thank Dr. Biafra Ahanonu for his assistance in data processing and Dr. P. Jeffrey Conn for providing VU0467154.

## Author information

Seongsik Yun, Ben Yang, Madison M. Martin, Nai-Hsing Yeh, Anis Contractor, & Jones G. Parker

## Contributions

S.Y. performed all imaging, behavior experiments, and histological experiments. B.Y. performed surgeries and assisted with imaging experiments. M.M.M. performed mouse surgeries and oversaw mouse breeding. N-H.Y. & A.C. conducted electrophysiology experiments. S.Y. and J.G.P. designed all experiments, performed all data analysis, and wrote the manuscript with input from the co-authors.

## Ethics declarations

The authors declare no competing interests.

## Supplementary Information

**Extended Data Fig. 1:**
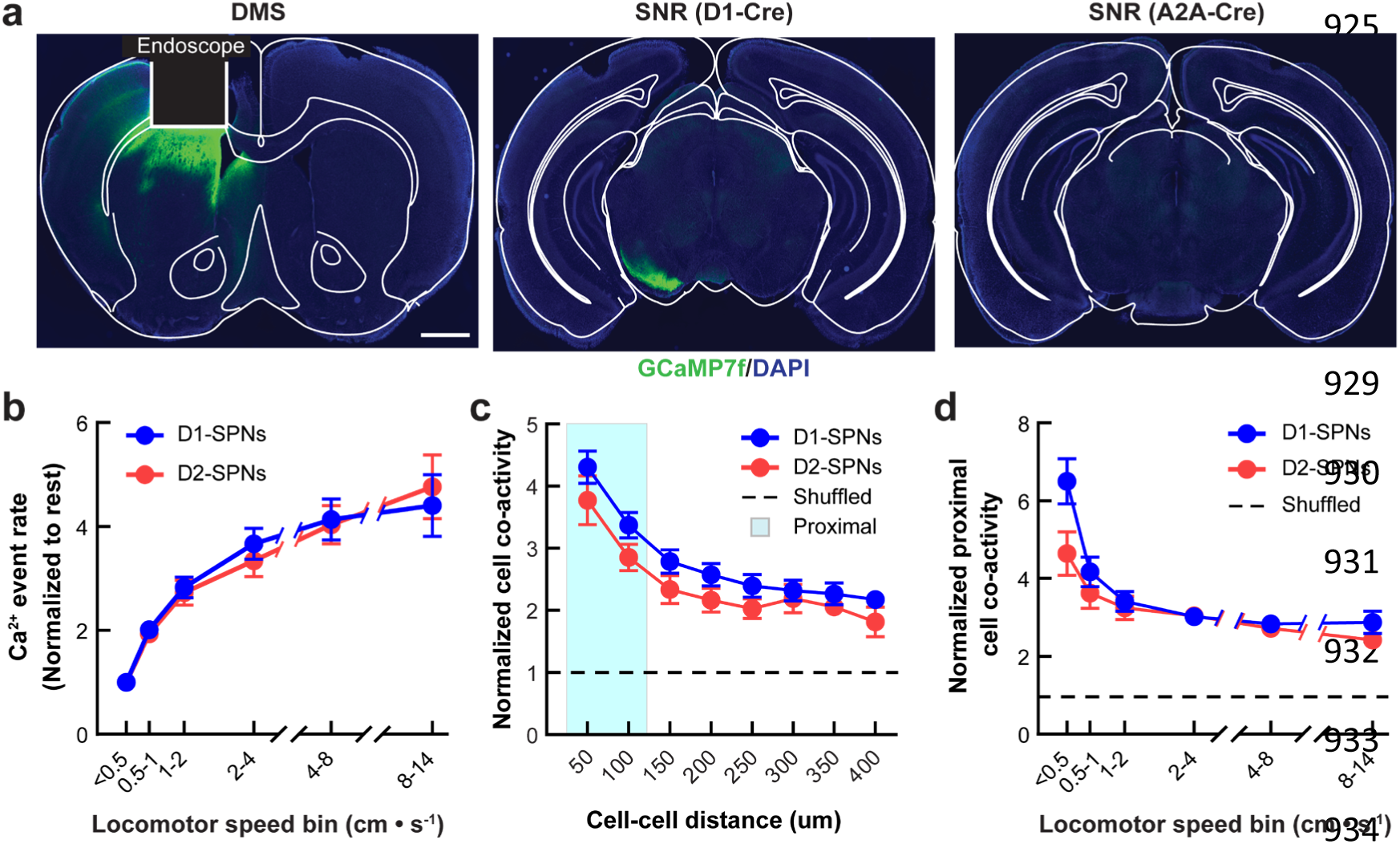
Quantifying normal D1- and D2-SPN ensemble dynamics. **a**, Representative coronal brain sections of dorsomedial striatum and substantia nigra from experimental D1- or A2A-Cre mice (*green*: anti-GFP; *blue*: DAPI nuclear stain; scale bar: 1 mm). White lines indicate the position of the implanted microendoscope and boundaries of brain areas. **b**, Ca^2+^ event rates in D1- and D2-SPNs at different speed bins normalized to event rate levels at rest (locomotor speed < 0.5 cm·s^-1^). **c**, Co-activity (jaccard index) of D1-SPN or D2-SPN pairs during movement (locomotor speed >= 0.5 cm·s^-1^) versus the separation of cell pairs normalized to temporally shuffled datasets (*dashed line*). Cyan shading indicates proximally (25–125 μm) cell pairs. **d**, Co-activity of proximal D1- and D2-SPN pairs at increasing bins of locomotor speed, normalized to temporally shuffled comparisons (*dashed line*). Data are expressed as mean ±s.e.m. (*N* = 11 D1-Cre and *N* = 10 for A2A-Cre mice; data were averaged across all recordings following vehicle only treatment).

**Extended Data Fig. 2:**
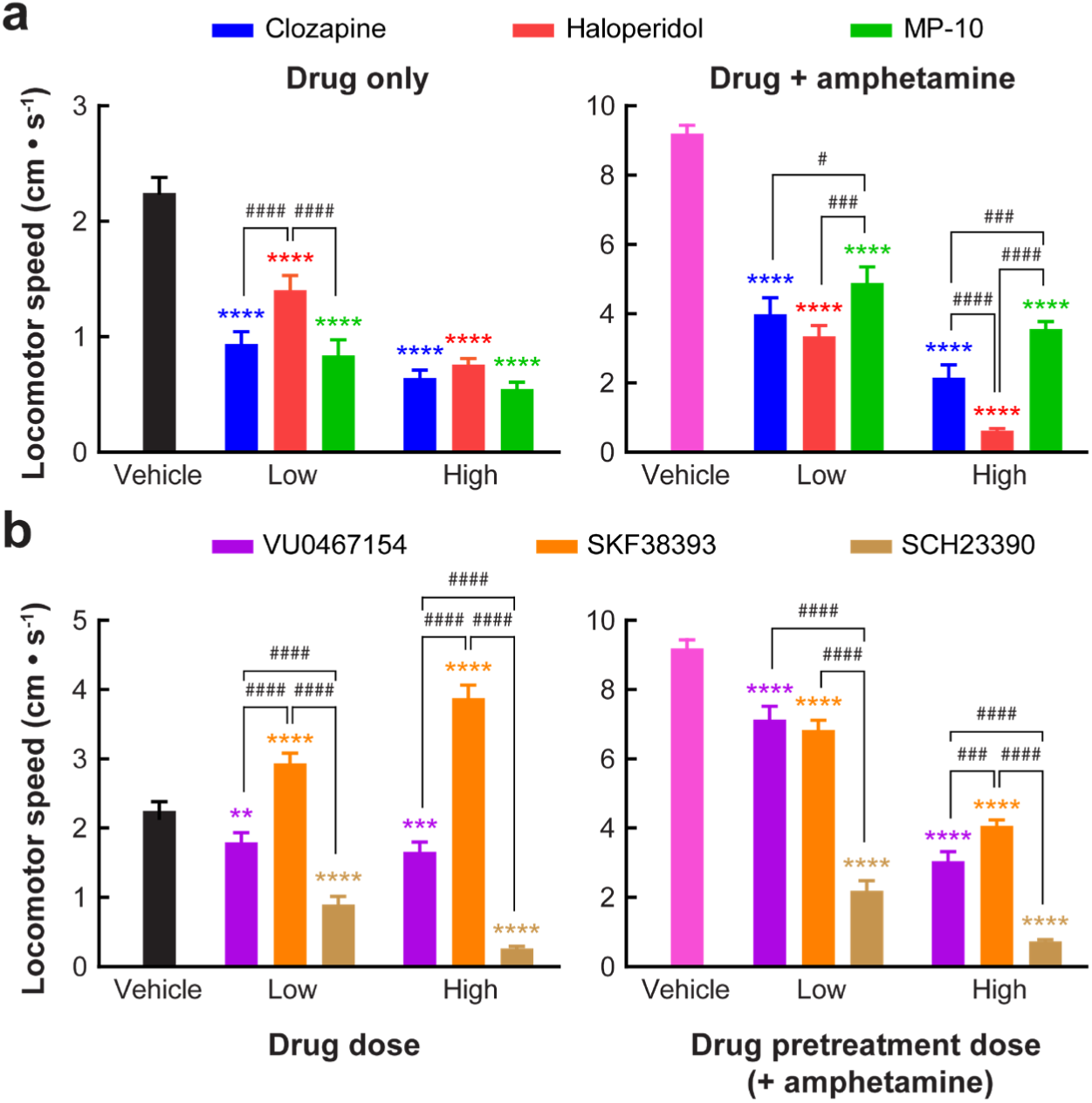
Effects of antipsychotic and D1-SPN-targeted drugs on normal and amphetamine-driven locomotor activity. **a**, **b**, Running speed following treatment with vehicle, antipsychotics (**a**) or D1-SPN-targeted compounds (**b**) with (*right*) or without amphetamine co-treatment (*left*). Data are represented as mean ± s.e.m. (*N* = 31 mice; \*\*\*\**P* < 0.0001, \*\*\**P* < 0.001, \*\**P* < 0.01 and \**P* < 0.05 compared to vehicle only treatment; ^####^*P* < 0.0001, ^###^*P* < 0.001 and ^#^*P* < 0.05 comparing the different drug treatment combinations; Holm-Sidak’s multiple comparison test).

**Extended Data Fig. 3:**
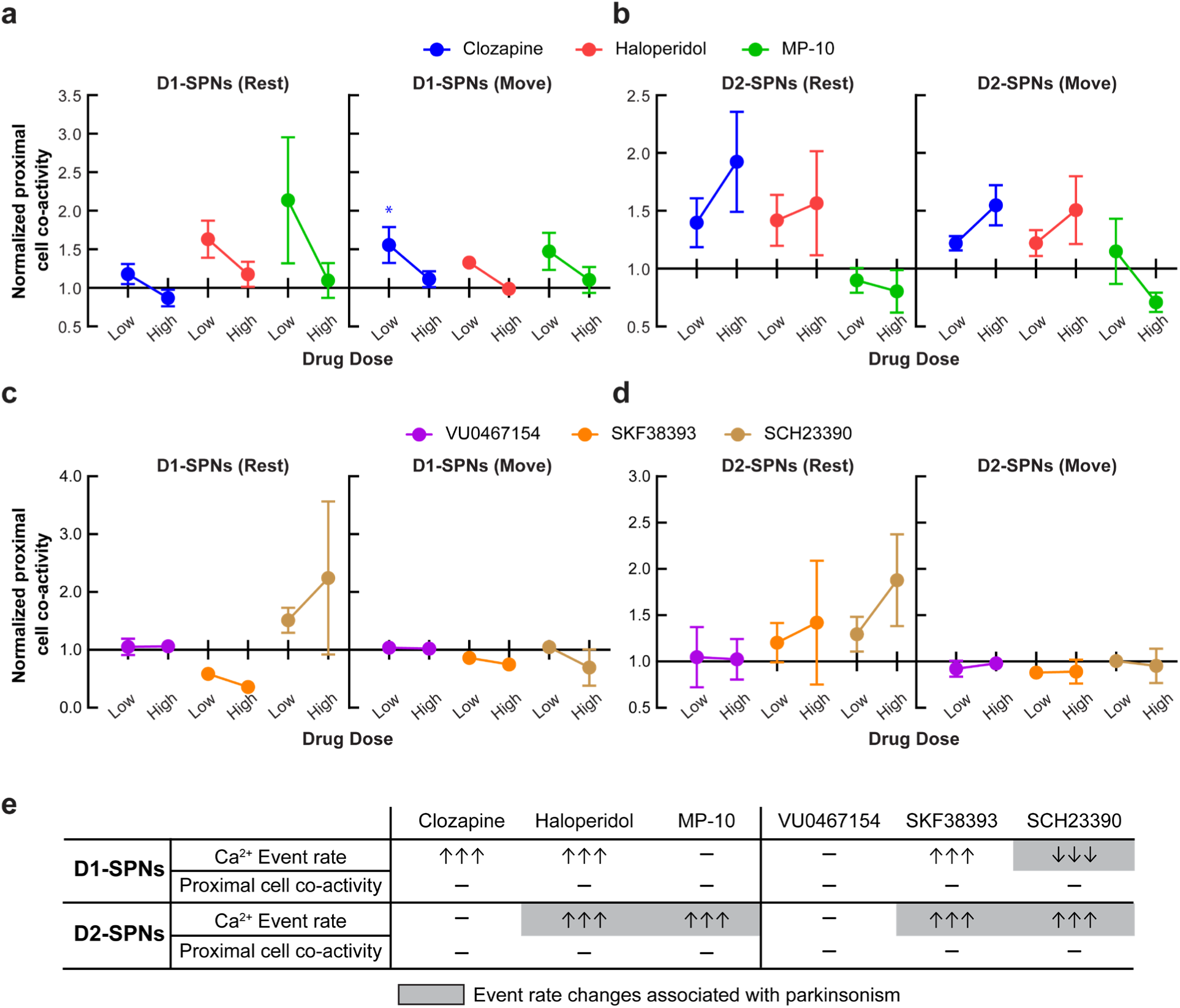
Effects of antipsychotic and D1-SPN-targeted drugs on the proximal co-activity of D1- and D2-SPN under normal conditions. **a**, **b**, Co-activity of proximal D1- (**a**) and D2-SPN (**b**) pairs during rest and movement following clozapine, haloperidol, or MP-10 administration. **c**, **d**, Proximal co-activity in D1- (**c**) and D2-SPNs (**d**) during rest and movement following VU0467154, SKF38393, or SCH23390 administration. Proximal co-activity values were first binned by locomotor speed, normalized to comparisons in temporally shuffled datasets within each speed bin, and normalized to the corresponding values following vehicle only treatment. Data are expressed as mean ± s.e.m. (*N* = 11 D1-Cre and *N* = 10 A2A-Cre mice; \**P* < 0.05 compared to vehicle treatment; Holm-Sidak’s multiple comparison test). **e**, Summary of the effects of different antipsychotic and D1-SPN-targeted drugs on D1- and D2-SPN ensemble dynamics under normal conditions. Each arrow denotes mean changes of >= 50% at the highest dose tested (see also Fig. 2c, d).

**Extended Data Fig. 4:**
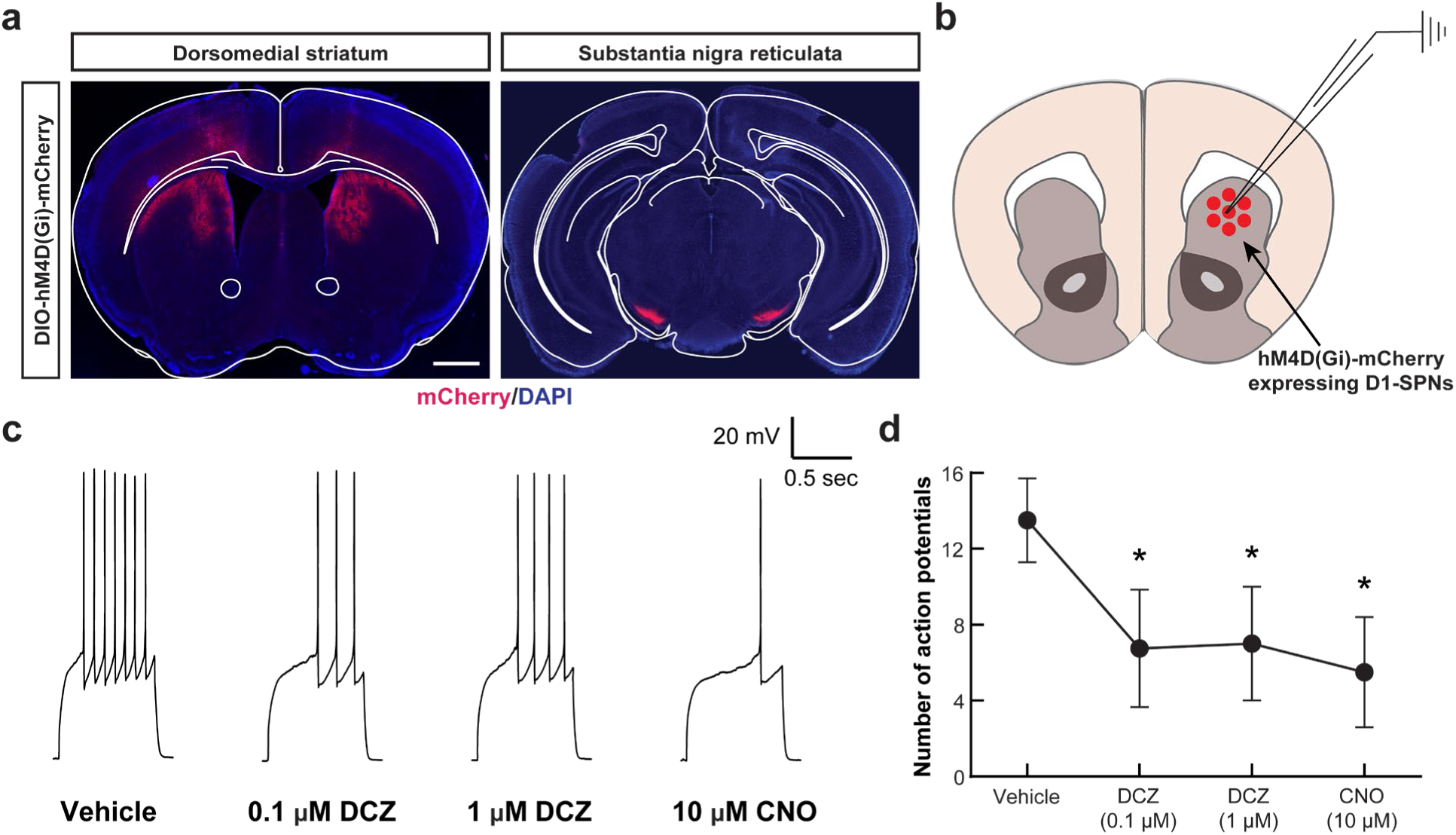
Histological and electrophysiological characterization of hM4Di-mCherry expression and function. **a**, Representative coronal brain sections of dorsomedial striatum and substantia nigra from experimental D1-Cre mice. Red indicates hM4Di-mCherry and blue indicates DAPI nuclear stain. Scale bar, 1 mm (see also Fig. 4). **b**, We performed patch-clamp electrophysiological recordings from hM4Di-mCherry-expressing neurons in the DMS of D1-Cre mice. **c**, Representative traces of action potential responses to 250 pA current injection. **d**, Number of action potentials following vehicle, DCZ or CNO treatment. Data are expressed as mean ± s.e.m. (*N* = 4; \**P* < 0.05 compared to vehicle treatment; Holm-Sidak’s multiple comparison test).

**Supplementary Table 1.**
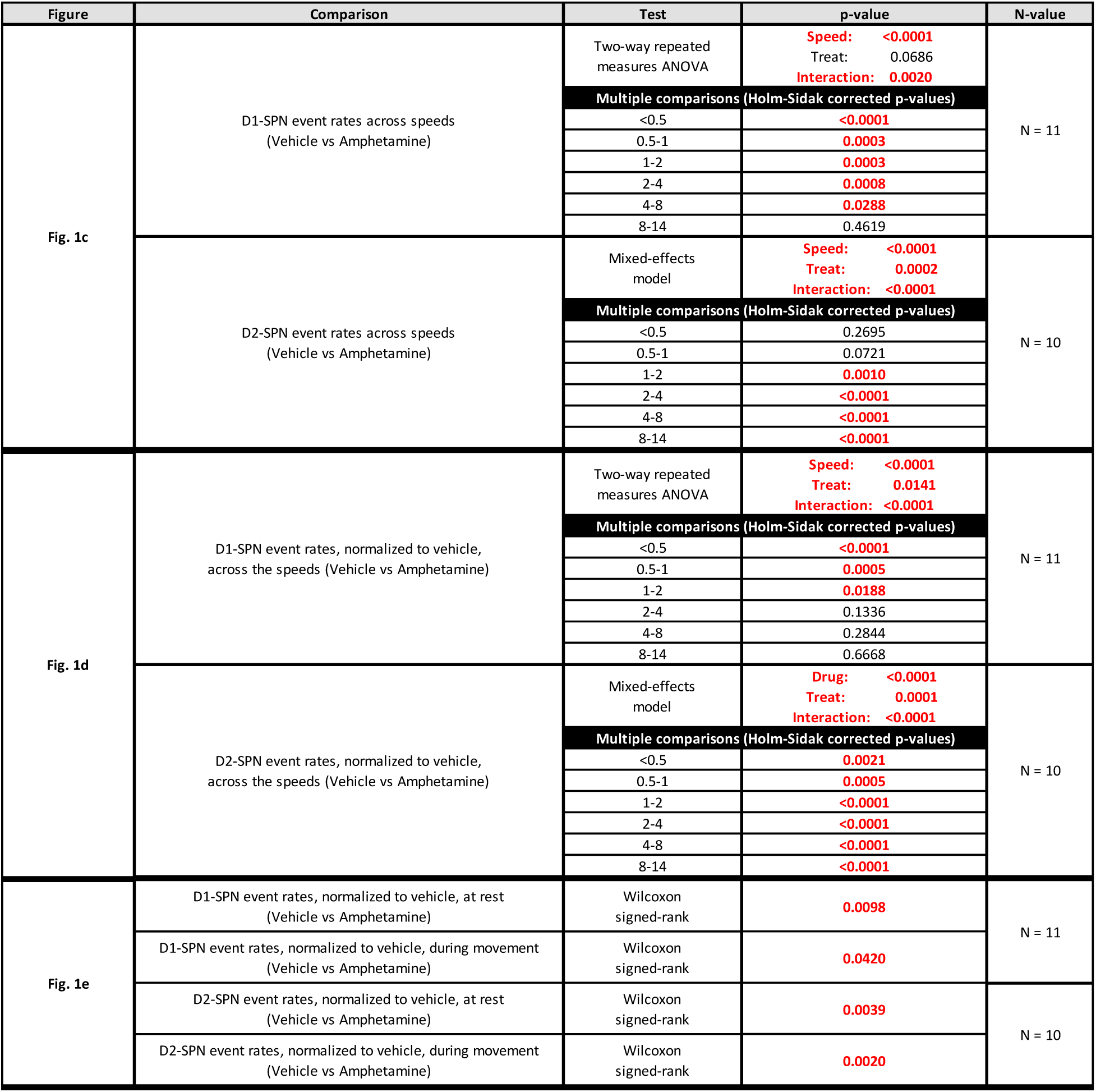
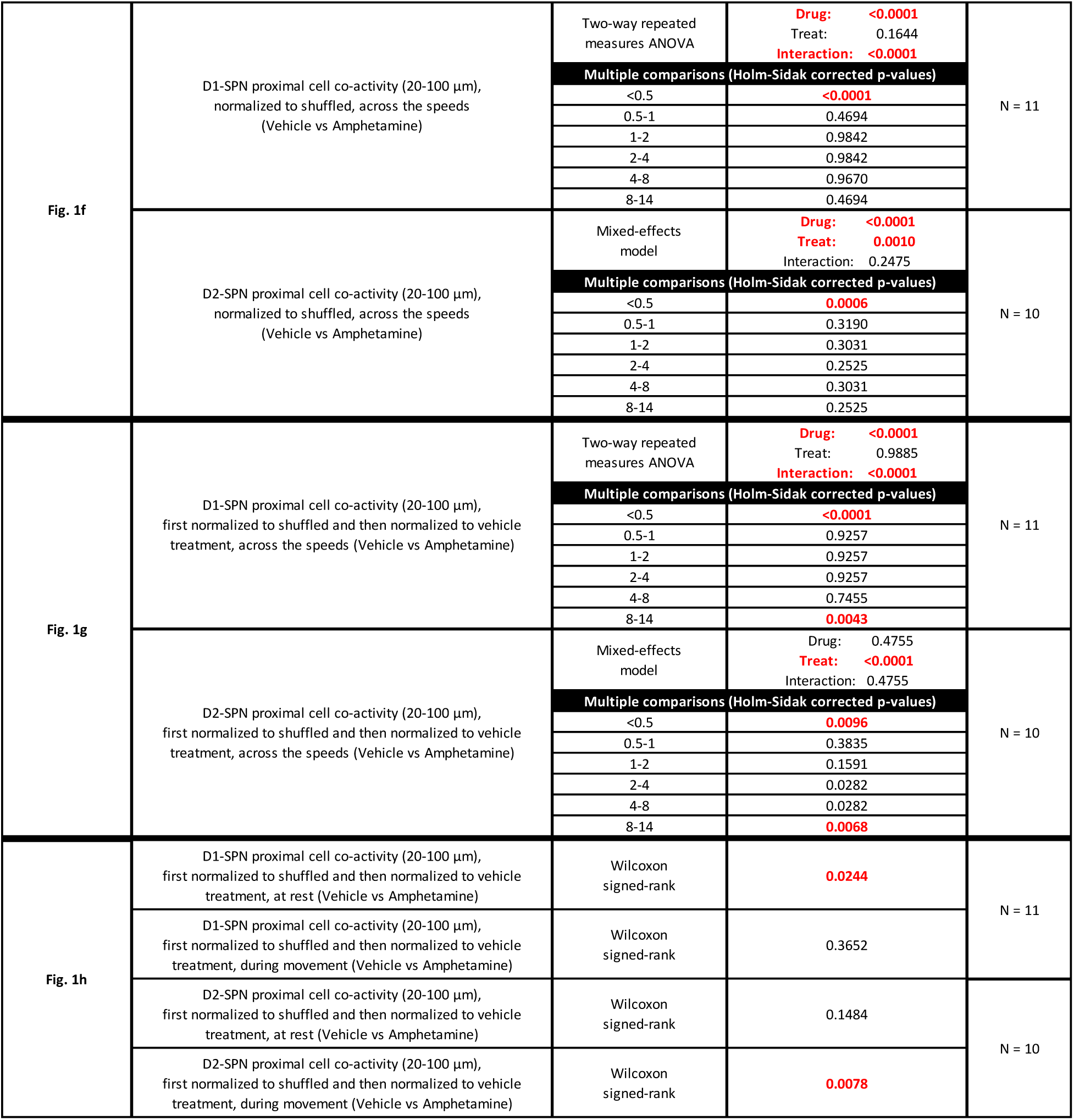
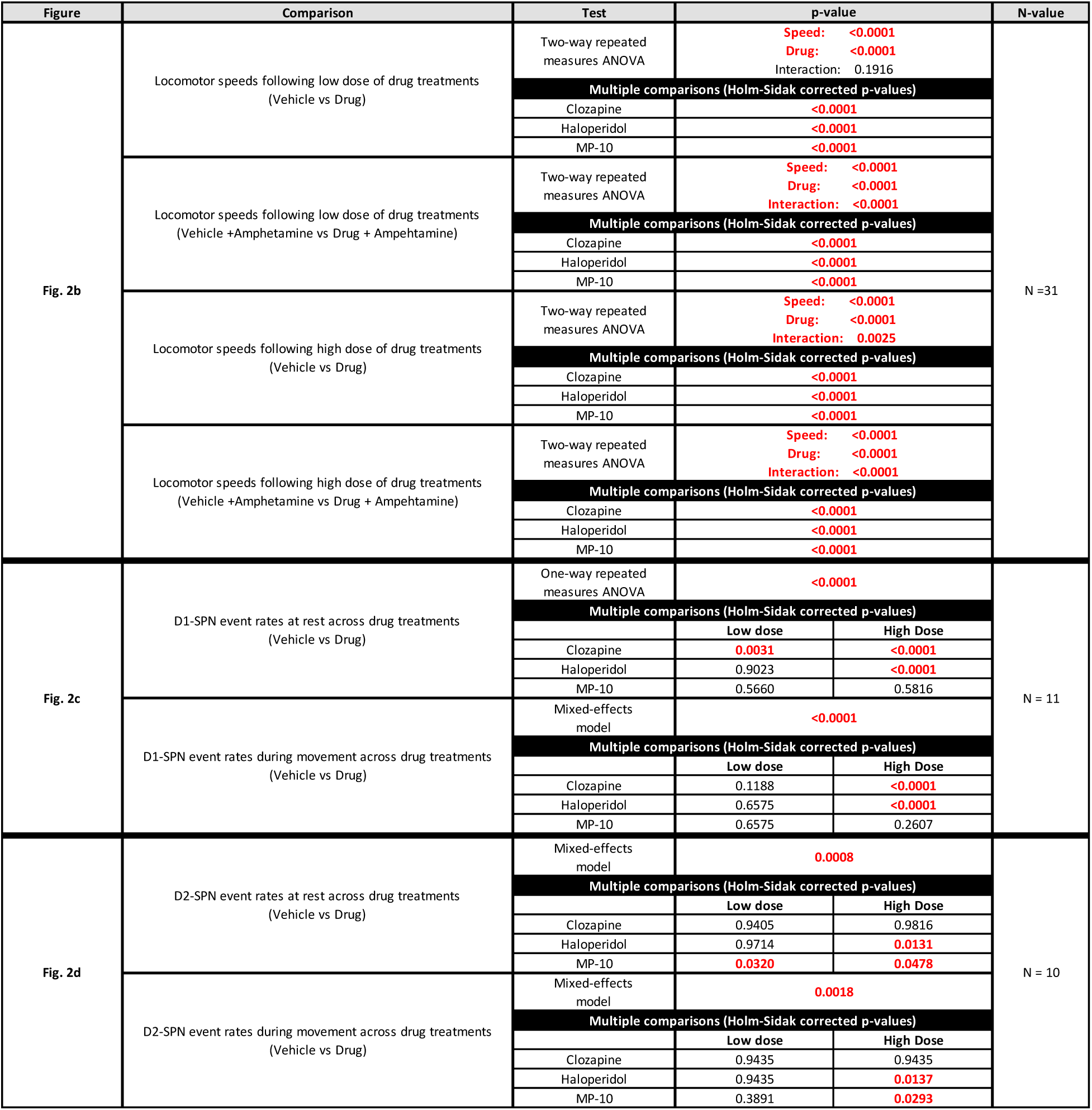
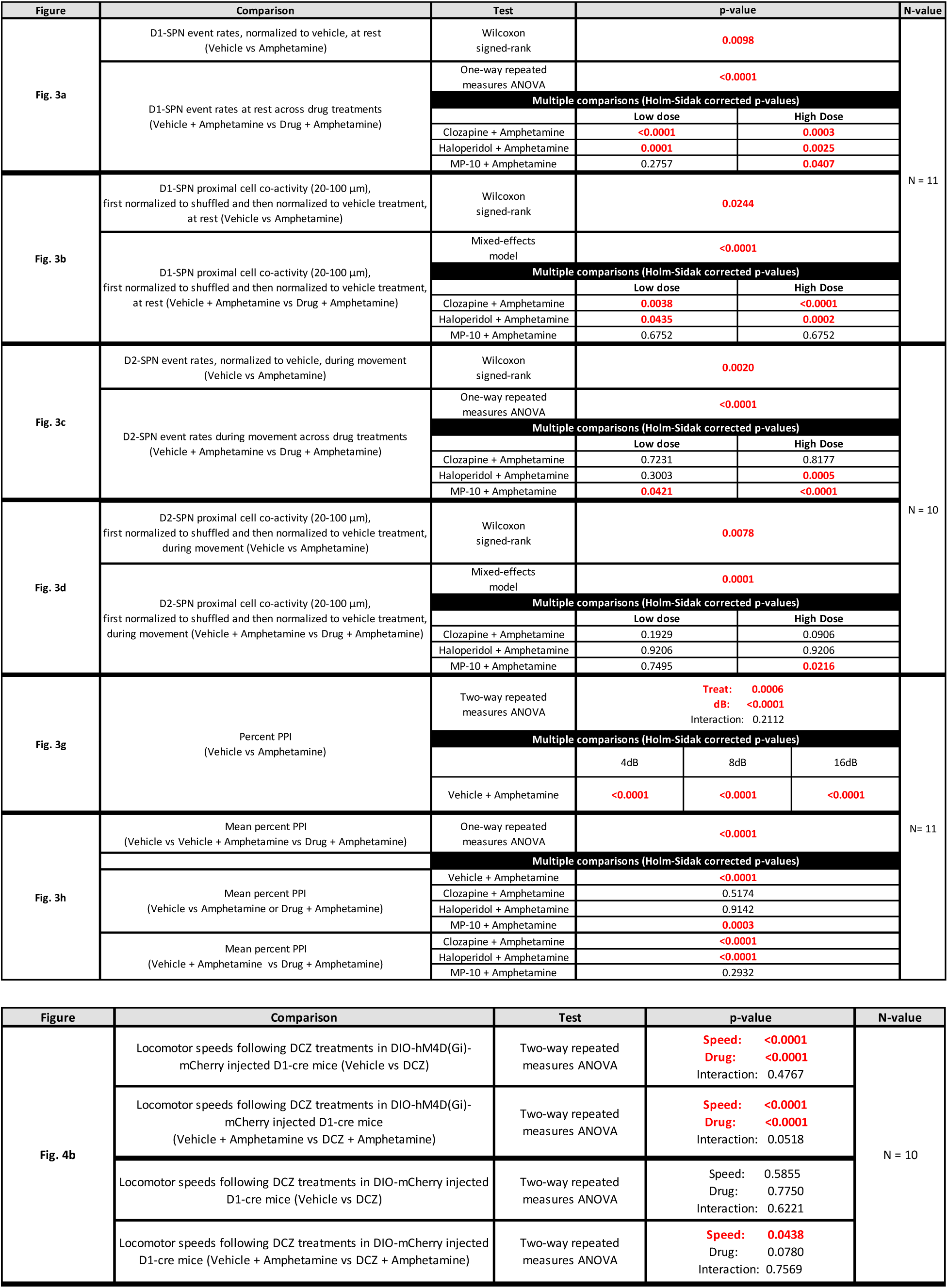
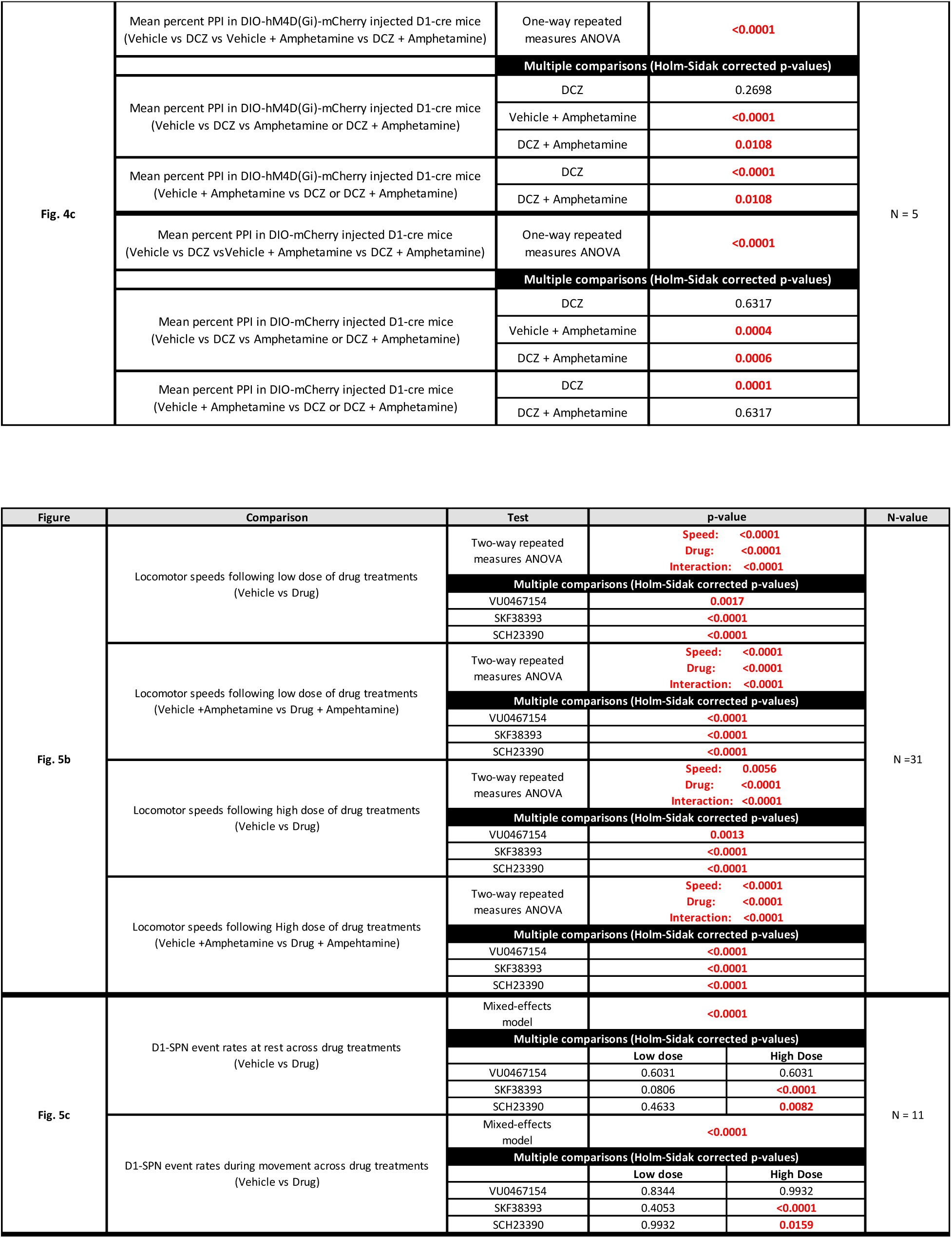
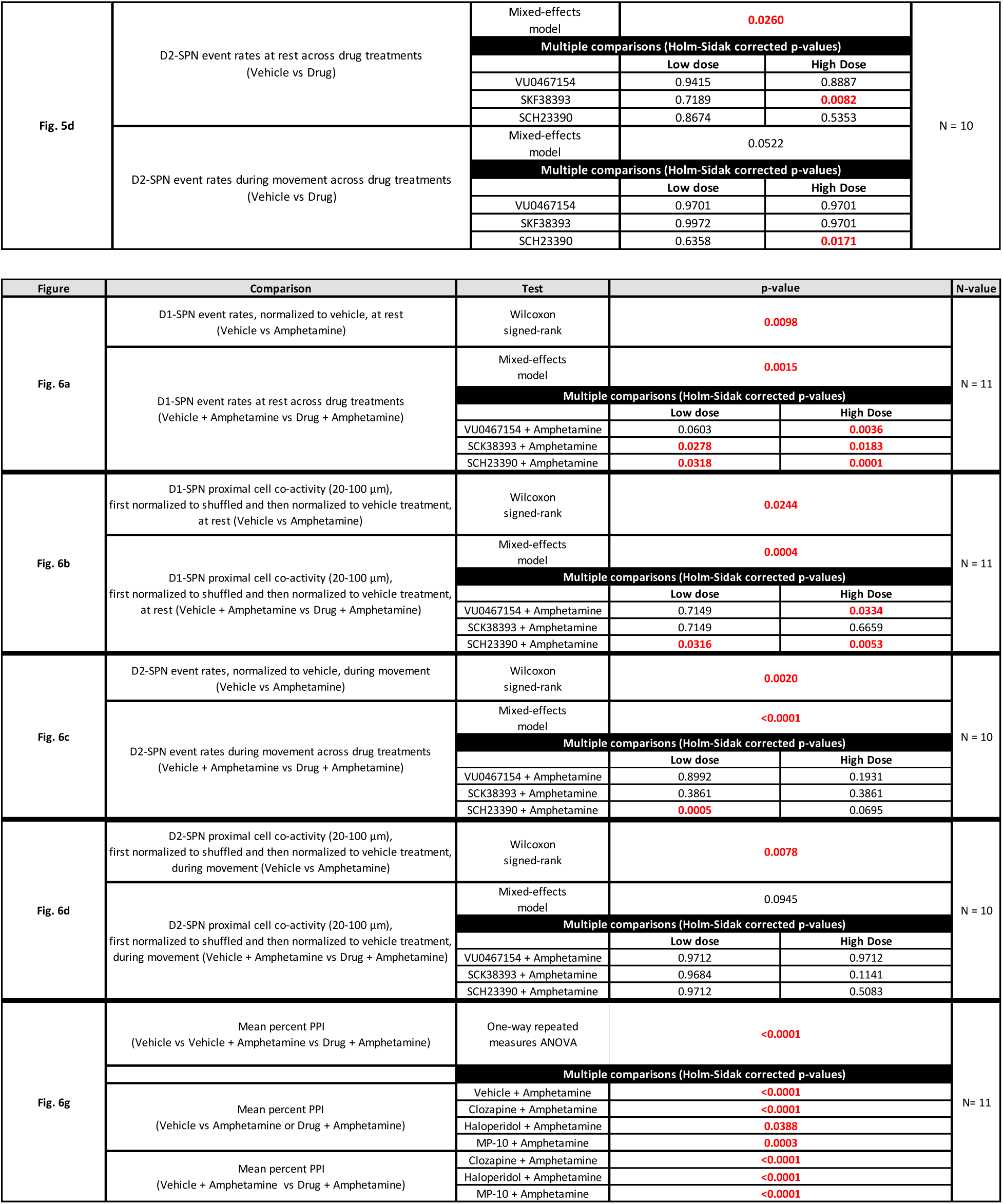
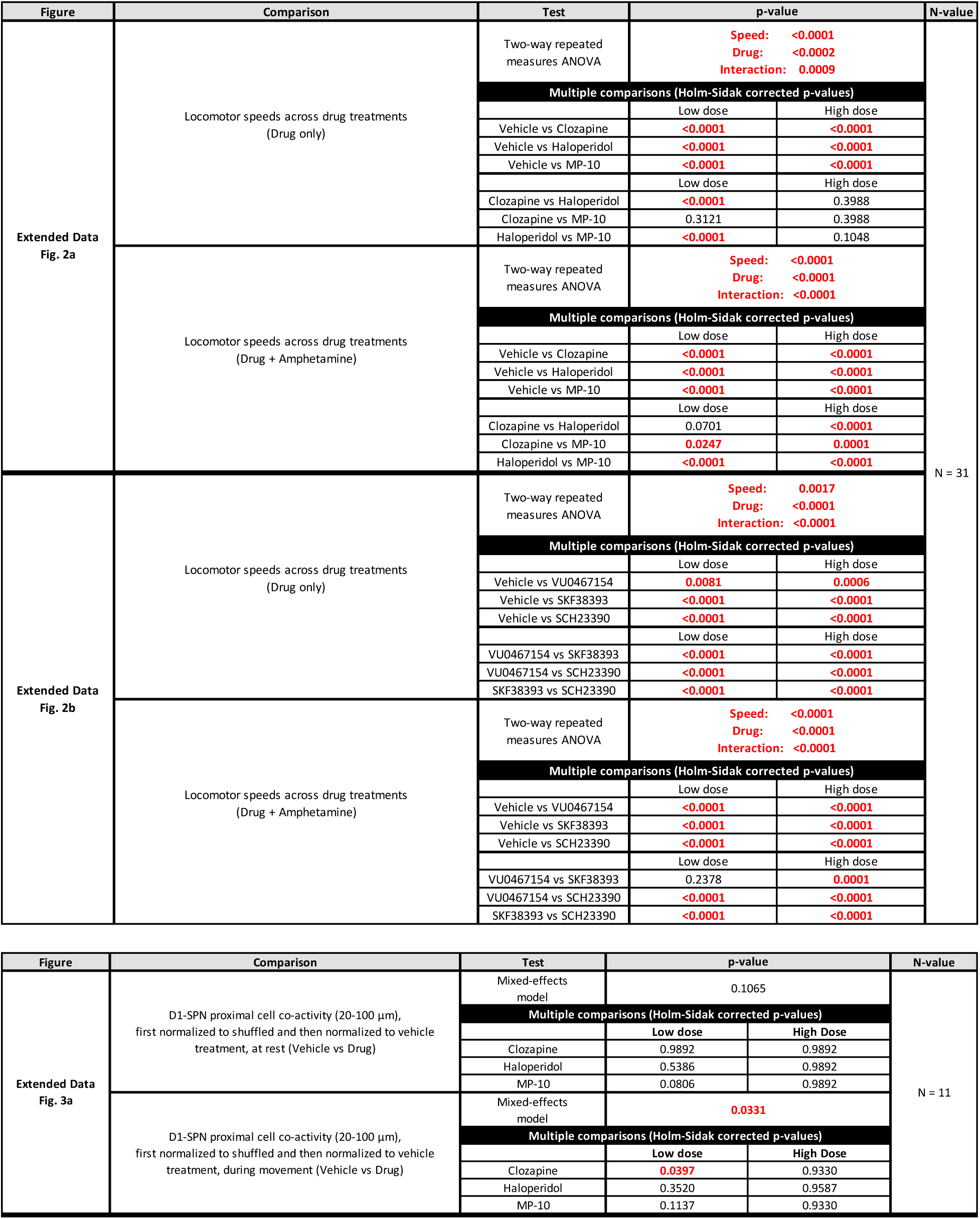
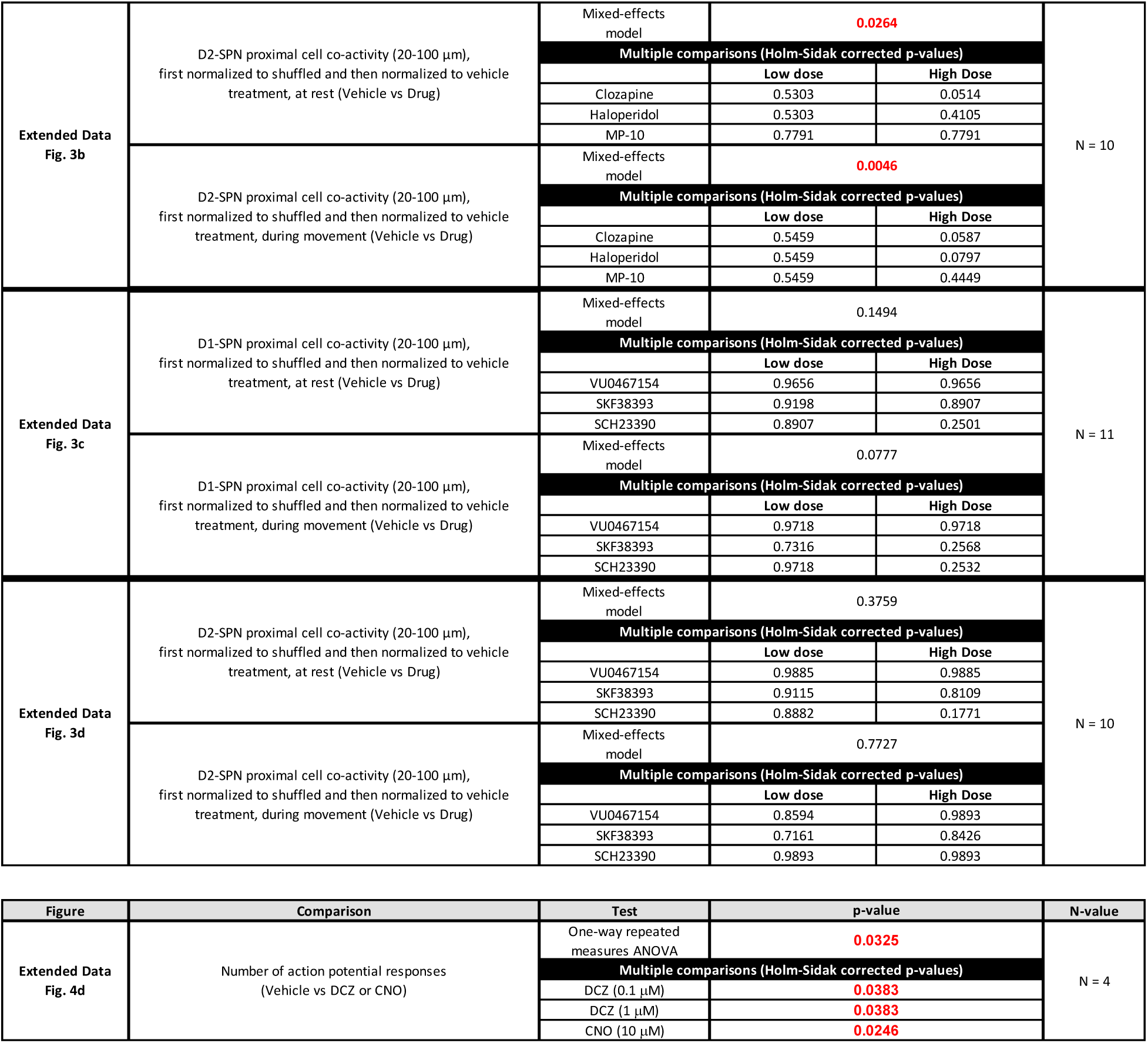

